# Gut-derived ammonia modulates hypothalamic stress responses during viral infection

**DOI:** 10.64898/2026.07.06.736796

**Authors:** Henrique G. Colaço, Anna Gundacker, Aubrey Burrett, Christian Grozev, Anna Hofmann, Joana Séneca, Lukas Endler, Joel Wong, Juan Sanchez, Maximilian Baumgartner, Christopher W. Fell, Alexander Lercher, Magdalena Siller, Zsofia Keszei, Csilla Viczenczova, Felix C. Richter, Yee Kwan Law, Laura Antonio-Herrera, Bethany Dearlove, Lorenz Balcar, Georg Kramer, Thomas Reiberger, Petra Pjevac, Clarissa Campbell, Daniela D. Pollak, Andreas Bergthaler

## Abstract

The gut–brain axis integrates microbial and host metabolism to regulate systemic physiology, yet its role during viral infection remains poorly defined. Viral infection induces behavioral changes and neuroendocrine stress responses accompanied by profound alterations in gut microbial metabolism. Here, we show that chronic viral infection in mice increases systemic levels of microbiota-derived ammonia in a CD8⁺ T cell–dependent manner. Increased ammonia accumulates in the brain and selectively activates neurons within the paraventricular hypothalamus (PVH), driving corticosterone release into the circulation. Pharmacological inhibition of ammonia detoxification exacerbates these effects, leading to increased corticosterone levels, aggravated sickness behavior, and dampened antiviral responses. Together, these findings identify gut-derived ammonia as a previously unrecognized immunometabolic signal linking antiviral T cell responses to hypothalamic control of systemic stress during viral infection.

**Graphical abstract:** 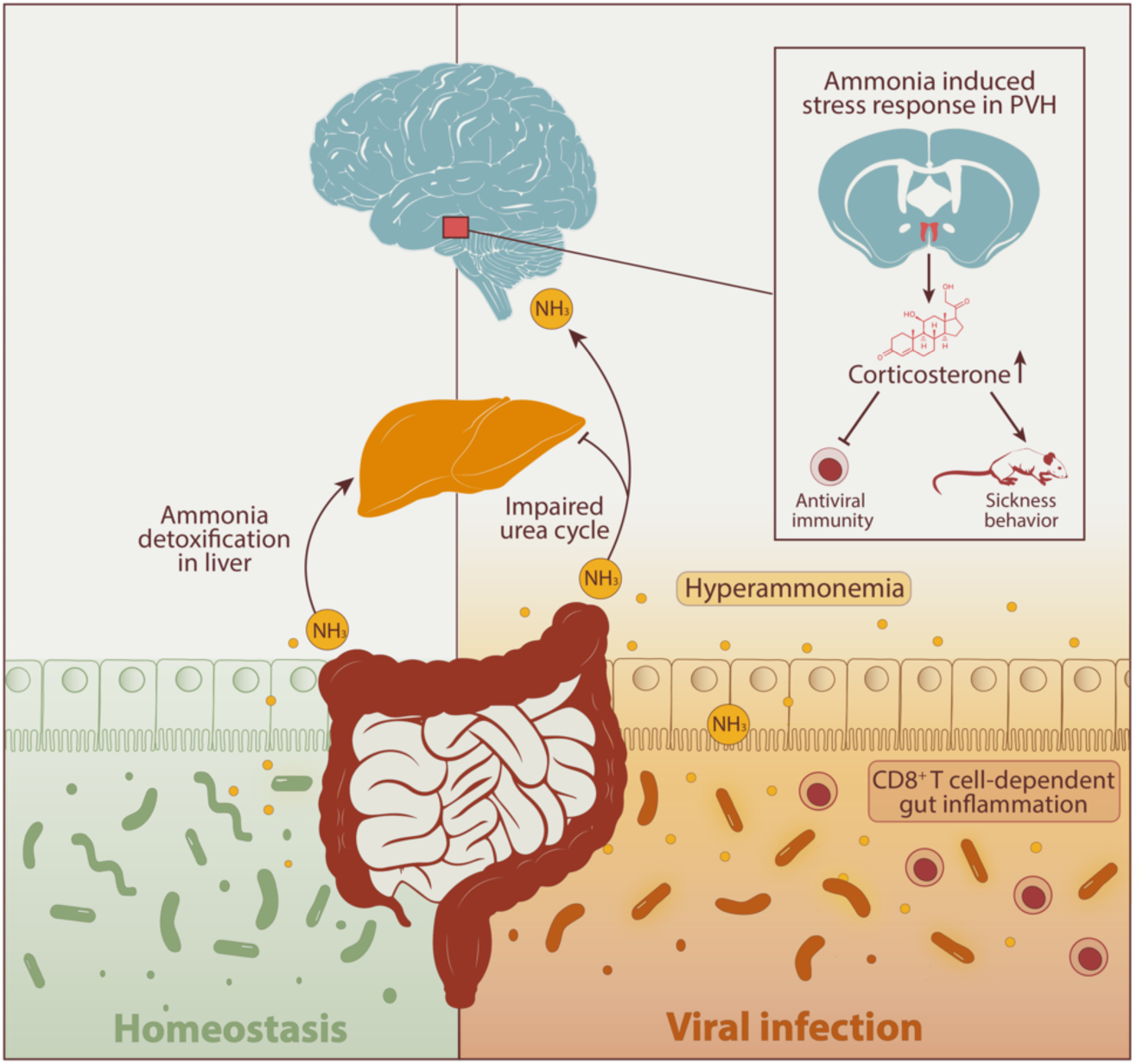

## Introduction

Infections rewire whole-body circuits and impose drastic immune, behavioral and metabolic changes to the host^1,2^. The central nervous system plays a crucial role both at sensing the danger of infection and orchestrating appropriate systemic adaptations^3,4^. Inflammatory responses in the periphery relay information to the brain either by stimulating the autonomic nervous system (e.g., through vagal afferents) or through cytokine signaling via the blood-brain barrier^3,5^. Following such stimuli, the brain coordinates systemic adaptations including the stress response system through the activation of the hypothalamus-pituitary-adrenal axis^6^. This is accompanied by a combination of evolutionarily conserved behavioral changes such as anorexia, social isolation and reduced libido, collectively known as sickness behavior^7^. Recent research has revealed multiple neuronal pathways activated during infection, including in the hypothalamus^8^ and brain stem^9^, highlighting the complex interaction between stimuli, neurocircuitry and outputs that govern systemic adaptations to infection.

Gut microbiota produce metabolites that affect numerous organismal functions^10^. Changes in gut microbiota composition have been reported in mouse models of viral infection, including lymphocytic choriomeningitis virus (LCMV)^11^ or influenza A virus^12^. Microbial metabolites originating in the gut have the potential to reach the brain^13^ and modulate physiological functions such as body temperature and feeding behavior in healthy mice^14^. Furthermore, several associations between the gut microbiota and brain function have been described in the context of neurodegeneration, autism and mood disorders^15^. Nevertheless, the role of microbe-derived metabolites in modulating brain function, stress responses, and sickness behavior during infection remain elusive.

Among microbially derived metabolites, small gaseous molecules such as ammonia, hydrogen sulfide, and nitric oxide have generated particular interest due to their capacity to diffuse throughout the body and exert multiple signaling functions^16–19^. Ammonia is produced as an end-product of amino acid catabolism and is therefore involved in several metabolic pathways across tissues like the gut, liver, kidney and brain^20^. Despite being typically considered a harmful waste product, due to its neurotoxic effect, for example in patients with liver disease^21^, signaling properties of ammonia in other contexts have just begun to be understood^22^.

In this study, we report a previously unrecognized role of ammonia as a messenger between the gut and the brain during viral infection. We observed that T cell-mediated systemic inflammation in mice triggers an increase in blood ammonia that is dependent on the gut microbiota. Gut-derived ammonia reaches the brain in high amounts during infection and stimulates the paraventricular hypothalamus (PVH) leading to the release of glucocorticoids into circulation, which affect behavioral and immune responses to infection. Our work underlines a previously unrecognized communication axis involving T cells, the gut microbiota and the brain, and highlights the importance of peripheral inputs in regulating glucocorticoid production during infection.

## Results

### Chronic LCMV infection causes hyperammonemia in a CD8^+^ T cell-dependent manner

We previously reported that chronic infection with LCMV Clone 13 (Cl13) leads to an increase in circulating ammonia at 8 days post-infection (dpi)^23^. To explore the kinetics of infection-induced hyperammonemia and its correlation with disease progression, we sampled mouse blood longitudinally during infection. We observed a steady increase of blood ammonia during the first week of infection with a maximum at 8 dpi (Figure 1a), which coincides with the peak of the adaptive immune response and associated metabolic and behavioral changes^24^. Notably, despite the prolonged weight loss and viral persistence associated with LCMV Cl13 infection, hyperammonemia is resolved within 12 dpi and ammonia levels remain low for over six weeks (Figure 1a).

**Figure 1:**
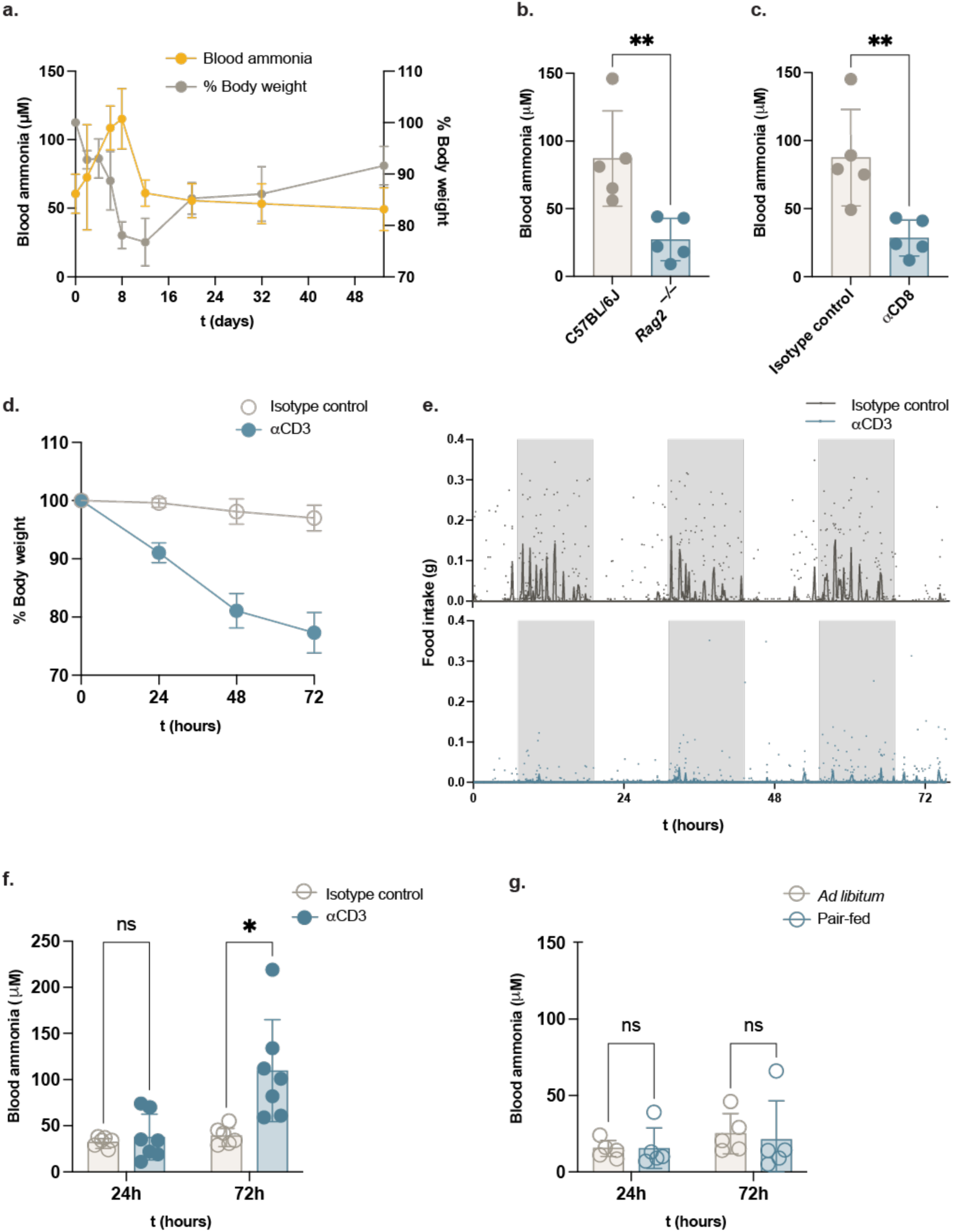
Blood ammonia increases during LCMV infection in a CD8 T cell-dependent manner. (a) Kinetics of blood ammonia and body weight progression in C57BL/6J mice infected intravenously with 2×10^6^ focus forming units LCMV Cl13. Data represents mean±SD, n=5 mice per group with longitudinal measurements. (b) and (c) Blood ammonia measurements at 8 days post LCMV Cl13 infection in *Rag2^−/−^*mice (b) or C57BL/6J pre-treated with an anti-CD8a antibody or an isotype control (c). Data shows mean±SD, n=5 mice per group. Statistical analysis: unpaired t-test. (d) Kinetics of body weight progression in C57BL/6J mice treated with 50 μg anti-CD3 (to induce sterile inflammation) or isotype control. Data represents mean±SD, n=6-7 mice per group, pooled from two independent experiments. (e) Food intake recorded in metabolic cages in C57BL/6J mice treated with 50 μg anti-CD3 or isotype control. Each point represents one mouse, lines indicate the median of each group and shaded areas indicate the dark cycle. n=4 mice per group. (f) Kinetics of blood ammonia in C57BL/6J mice treated with 50 μg anti-CD3 or isotype control. Data represents mean±SD, n=6-7 mice per group, pooled from two independent experiments. Statistical analysis: multiple unpaired t-tests with Welch correction. (g) Blood ammonia measurements at 24 h and 72 h of *ad libitum* feeding or pair feeding. Pair-fed mice were given the same amount of food as consumed by αCD3-treated mice, as shown in (e). Data shows mean±SD, n=5 mice per group. Statistical analysis: multiple unpaired t-tests with Welch correction.

The peak of hyperammonemia at 8 dpi suggests an involvement of the adaptive immune response. To test this hypothesis, we infected *Rag2*^−/−^ mice, which lack mature T and B cells, with LCMV Cl13 and observed lower levels of blood ammonia at 8 dpi compared to wildtype controls (Figure 1b). As the LCMV infection model heavily relies on cytotoxic CD8^+^ T cells, we employed an antibody treatment to deplete CD8a^+^ cells from the circulation of wildtype mice (Supplementary figure 1a) and observed that depleted mice show decreased blood ammonia at 8 dpi (Figure 1c). Treatment of mice with an antibody against chemokine receptor type 3 (CXCR3), which mediates CD8^+^ T cell activation and proliferation ^25,26^, caused a decrease in effector CD44^+^ and in virus-specific GP33^+^ splenic T cells (Supplementary figure 1b), and partially rescued mice from hyperammonemia (Supplementary figure 1c). Conversely, genetic ablation of type 1 interferon signaling, which is essential for the innate immune response in the first days of infection, caused no changes in blood ammonia at 8 dpi (Supplementary figure 1d). Together, this data shows that the adaptive immune response, particularly the activation of cytotoxic T cells, is necessary for infection-induced hyperammonemia.

Based on these results, we reasoned that T cell activation might be sufficient to increase blood ammonia even in the absence of viral infection. To test this, we injected an anti-CD3 antibody as a model of sterile T cell activation and immunopathology^27,28^. Anti-CD3 treated mice developed significant body weight loss (Figure 1d), recapitulating the severity of LCMV-induced immunopathology albeit in a shorter time frame. In-depth characterization of these mice using metabolic cages revealed signs of anorexia (Figure 1e), decreased movement (Supplementary figure 1e), along with changes in respiratory exchange ratio (RER) rhythms (Supplementary figure 1f), which also align with the sickness behavior and metabolic reprogramming observed in LCMV Cl13 infection^24,29^. In addition, anti-CD3 treated mice developed hyperammonemia within 72h (Figure 1f). In contrast, treatment with polyinosinic:polycytidylic acid [Poly(I:C)], a viral RNA analogue that stimulates TLR3 and triggers a T cell-independent systemic inflammation, had no effect on blood ammonia levels (Supplementary figure 1g). Together, our data show that antigen-independent T cell activation is sufficient to induce hyperammonemia and that viral sensing and type I interferon responses are dispensable for this effect.

Anorexia is a hallmark of infection and major driver of metabolic changes. To test if food restriction is sufficient to induce hyperammonemia in the absence of an inflammatory response, we fed naïve mice with the same amount of food consumed by anti-CD3 treated mice. We found that blood ammonia levels during pair-feeding were indistinguishable from mice fed ad-libitum (Figure 1g). The same applied to mice fed the same amount of food consumed by LCMV Cl13-infected mice (Supplementary figure 1h-i). In summary, our data suggests that T cell-mediated inflammation, but not food restriction, is sufficient to drive an increase in blood ammonia.

### Infection-induced hyperammonemia is dependent on the gut microbiota

LCMV Cl13 infection is characterized by widespread infiltration of CD8^+^ T cells in tissues, which causes immunopathology and metabolic rewiring of parenchymal cells. Several metabolic pathways result in production of ammonia as a byproduct, including amino acid deamination in the kidney, muscle and gut epithelium, and microbiota-dependent pathways in the gut. In parallel, ammonia is primarily eliminated in the liver through the urea cycle (Figure 2a). We therefore analyzed the contribution of different tissue-specific metabolic pathways to understand which is the major source of ammonia during infection. Expression of ammonia metabolism genes (Figure 2a) in the ileum and colon tissue revealed no changes during infection (Supplementary figure 2a-b). In the skeletal muscle, protein catabolism and adenosine monophosphate (AMP) deamination (via *Ampd1*) both release ammonia, which can significantly contribute to increasing its concentration in circulation^20,30^. However, transcriptional analysis of ammonia-generating genes in the skeletal muscle and kidney indicated a shutdown of ammonia production during infection (Supplementary figure 2c-d). This rendered increased amino acid catabolism in extra-hepatic tissues an unlikely explanation for hyperammonemia during infection.

**Figure 2:**
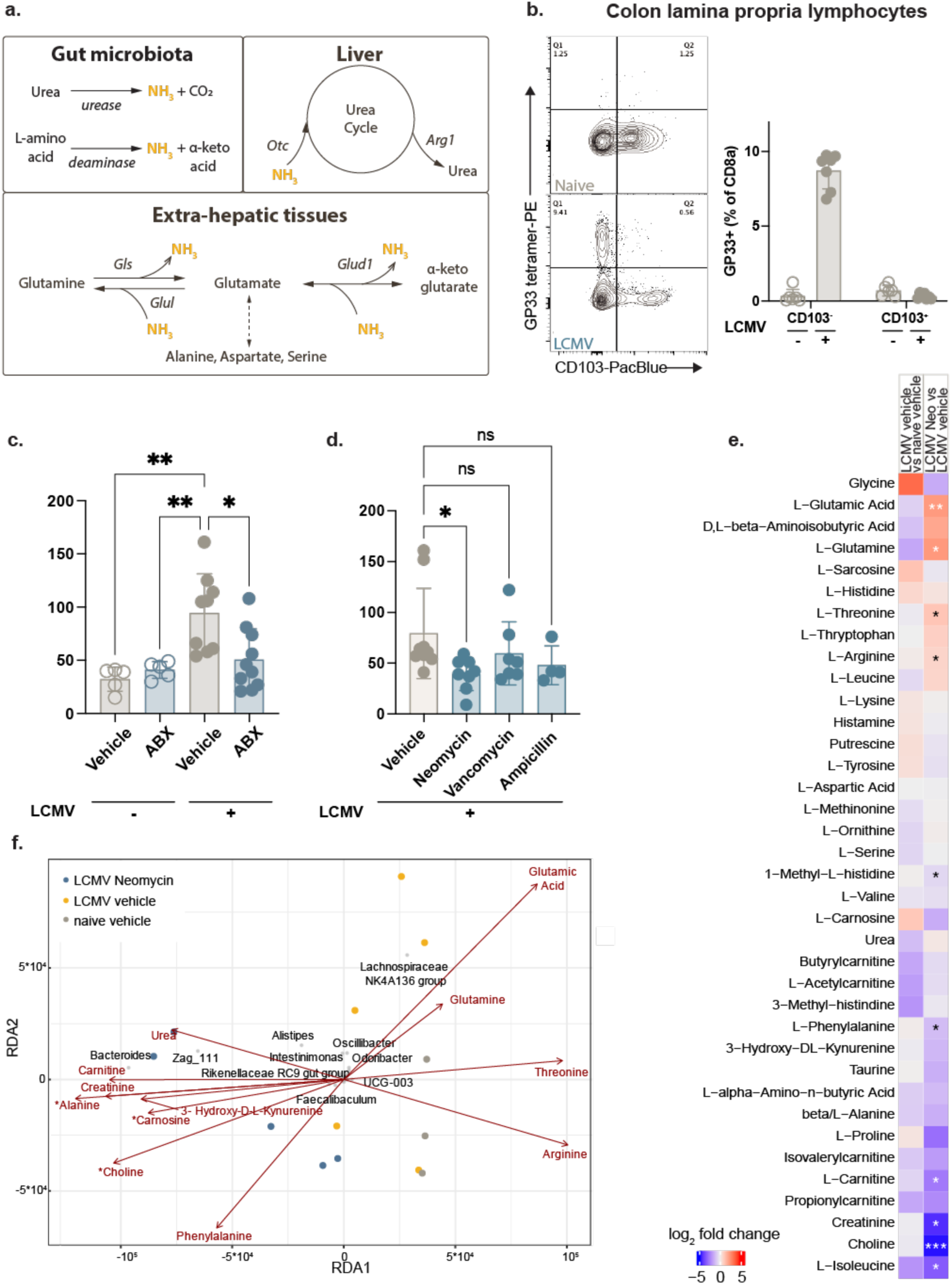
Hyperammonemia is dependent on the gut microbiota. (a) Summary of major metabolic pathways involving ammonia in the liver, gut microbiota and extra-hepatic tissues (skeletal muscle, kidney, intestinal epithelium and brain). (b) Flow cytometric analysis showing the percentage of virus-specific (GP33^+^) T cells that express the Trm T cell marker CD103 in lamina propria lymphocytes (LPL) in the colon of LCMV Cl13 infected mice at 8 dpi. Data shows mean±SD, n=6-7 mice per group pooled from two independent experiments. (c) Blood ammonia measurement at 8 days post LCMV Cl13 infection, and non-infected controls, of mice pre-treated with antibiotics (ABX) or sucralose (vehicle control). The large-spectrum antibiotic cocktail containing ampicillin, vancomycin, neomycin, metronidazole and gentamycin was given in the drinking water starting two weeks before infection and kept throughout the experiment. Data shows mean±SD, n=5-10 mice per group pooled from two independent experiments. Statistical analysis: one-way ANOVA with Tukey multiple comparisons test. (d) Blood ammonia measurement at 8 days post LCMV Cl13 infection in mice pre-treated with neomycin, vancomycin or ampicillin in the drinking water. Data shows mean±SD, pooled from two independent experiments, n=4-9 mice per group. Statistical analysis: one-way ANOVA with Tukey multiple comparisons test. (e) Metabolomics analysis of cecal samples from infected and neomycin-treated mice. Data represents the log2 fold change of the indicated comparisons, n=3-5 mice per group. Statistical analysis: unpaired t-test. (f) Redundancy analysis (RDA) biplot illustrating the relationship between genus-level microbial communities (dPCR-normalized) and selected metabolites. Samples are displayed as points colored by group and metabolites are represented as vectors indicating the direction and strength of their contribution to the ordination. The top 10 genera contributing most strongly to the separation are shown in black text. n=3-5 mice per group.

We then turned our attention to the gastrointestinal tract, as it is known as the major ammonia producing tissue in mammals^20^. T-cell dependent intestinal inflammation and dysbiosis have been described both in LCMV Cl13 infection^11,31,32^ and in the sterile inflammation model of anti-CD3 injection^28,33^. We observed an influx of GP33^+^ virus-specific T cells in the colon lamina propria at 8 days post LCMV Cl13 infection (Figure 2b), together with high expression of the pro-inflammatory cytokines IFNγ and TNFα (Supplementary figure 2e-f) and low expression of the tissue-resident memory T cell marker CD103^34^ (Figure 2b, Supplementary figure 2e-f). In parallel, we confirmed a CD8^+^ T cell-dependent increase in intestinal permeability in LCMV Cl13 infection, as judged by serum levels of lipopolysaccharide binding protein (LBP) (Supplementary figure 2g). We hypothesize that infiltration of circulatory T cells into the gut increases intestinal permeability and changes the gut microbial composition leading to an increase in blood ammonia. To test the contribution of the gut microbiota to infection-induced hyperammonemia, we treated mice with a cocktail of broad-spectrum antibiotics in the drinking water prior to LCMV Cl13 infection. We found a significant decrease in blood ammonia in infected, antibiotic-treated mice compared to infected, vehicle-treated controls (Figure 2c). Treatment with single antibiotics had distinct outcomes: neomycin, which primarily targets gram-negative bacteria, was effective in clearing blood ammonia whereas ampicillin and vancomycin, which mostly target gram-positive bacteria, had a negligible effect (Figure 2d).

Gut microbiota-derived ammonia originates from two major pathways: urea breakdown by urease-positive bacteria, and deamination of dietary protein by gut resident bacteria (Figure 2a)^35–37^. We measured urease enzymatic activity in fecal pellets 8 days after LCMV Cl13 infection and observed an increase in urease-dependent ammonia production during infection (Supplementary figure 2h). The effect was reverted when mice were pre-treated with an anti-CD8 depletion antibody (Supplementary figure 2h). We then employed two distinct small molecules, acetohydroxamic acid (AHA) and flurofamide (FFA), to inhibit urease activity during infection. Treatment with either drug resulted in a decrease in fecal urease activity but no difference in circulating ammonia levels (Supplementary figure 2i-j), indicating that urease activity is not sufficient to explain the increased pool of ammonia during infection.

To better understand the role of gut microbiota in driving hyperammonemia, we performed 16S rRNA gene amplicon sequencing of cecal contents of infected and neomycin-treated mice. In line with a previous study^11^, we found shifts in microbial composition upon LCMV Cl13 infection, including an increase in abundance of taxa affiliated with the genus *Bacteroides* (Supplementary figure 3a). Neomycin treatment during infection caused dramatic changes including a decrease in species richness and evenness (Supplementary figure 3b) accompanied by a significant difference in beta diversity between groups (Supplementary figure 3c). We complemented the analysis of these cecal contents with a targeted metabolomics panel focused on amino acids, amino acid derivatives and urea to better understand nitrogen balance and ammonia metabolism in the gut. Neomycin treatment during infection leads to a depletion of several amino acids including glutamine, glutamate, and arginine (Figure 2e). When comparing bacterial taxa and metabolites that are simultaneously enriched during infection and depleted upon neomycin treatment, we obtained a small set of potential ammonia-producing bacteria spanning two major phyla (Figure 2f, Supplementary figure 3a). The first, *Bacillota* (formerly known as *Firmicutes*), includes the *Clostridia* class, such as the *Lachnospiraceae* NK4A136 group and *Oscillibacter*. The second, *Bacteroidota* (formerly *Bacteroidetes*), is represented by the genera *Odoribacter* and *Alistipes*. Furthermore, changes in fecal glutamine and glutamate upon neomycin treatment correlated with abundance of the *Lachnospiraceae* NK4A136 group, *Oscillibacter* and *Odoribacter* (Figure 2f). *Clostridia* are generally known as effective amino acid degraders^38–40^, raising the possibility that amino acid fermentation by neomycin-sensitive gut bacteria increases ammonia output during infection. At the same time, *Bacteroidia* such as the *Alistipes* and *Odoribacter* genera are known for utilizing host-secreted proteins in conditions of limited dietary nitrogen influx^41^.

Blood ammonia levels result from a fine balance between host and microbiome-derived output from tissues and elimination in the hepatic urea cycle. We previously reported an impairment of urea cycle function during LCMV Cl13 infection^23^, which likely contributes to the systemic accumulation of ammonia. Nevertheless, infected, neomycin-treated mice sustain low levels of circulating ammonia despite presenting significant liver pathology, as judged by serum levels of alanine aminotransferase (ALT) (Supplementary figure 3d) and disrupted hepatic urea cycle, as judged by reduced transcript levels of *Otc* (Supplementary figure 3e). This data highlights the complexity of nitrogen balance during infection, in which both the production in the gut and deficient elimination in the liver contribute to the accumulation of ammonia in the blood.

Taken together, our data suggests that T cell-mediated intestinal inflammation causes a shift in microbial nitrogen metabolism in the gut, whereby multiple mechanisms including host-derived protein degradation and urease activity contribute to increased ammonia production, leading to hyperammonemia.

### Ammonia is readily distributed to the brain during infection

Due to its small size and capacity of crossing membranes both by diffusion and active transport^20,42^, ammonia can rapidly reach every tissue in the body. To understand how infection affects ammonia distribution and the metabolic flux of nitrogen, we employed a metabolite tracing approach based on systemic administration of heavy isotope labelled ^15^N-NH_4_Cl followed by LC-MS analysis in different tissues. The two metabolites with most pronounced nitrogen labelling were citrulline in the liver, which is compatible with ammonia incorporation in the urea cycle (Supplementary figure 4a), and glutamine in the brain, liver and kidney (Figure 3a-b). Labelling of other amino acids, such as glutamate, aspartate, arginine, serine, and alanine was marginal (Supplementary figure 4a), confirming previous reports showing that glutamine synthesis is the major route of ammonia detoxification in extra-hepatic tissues^43^. Interestingly, during LCMV infection we observed an increase in glutamine labelling in the brain but not in other tissues (Figure 3b).

**Figure 3:**
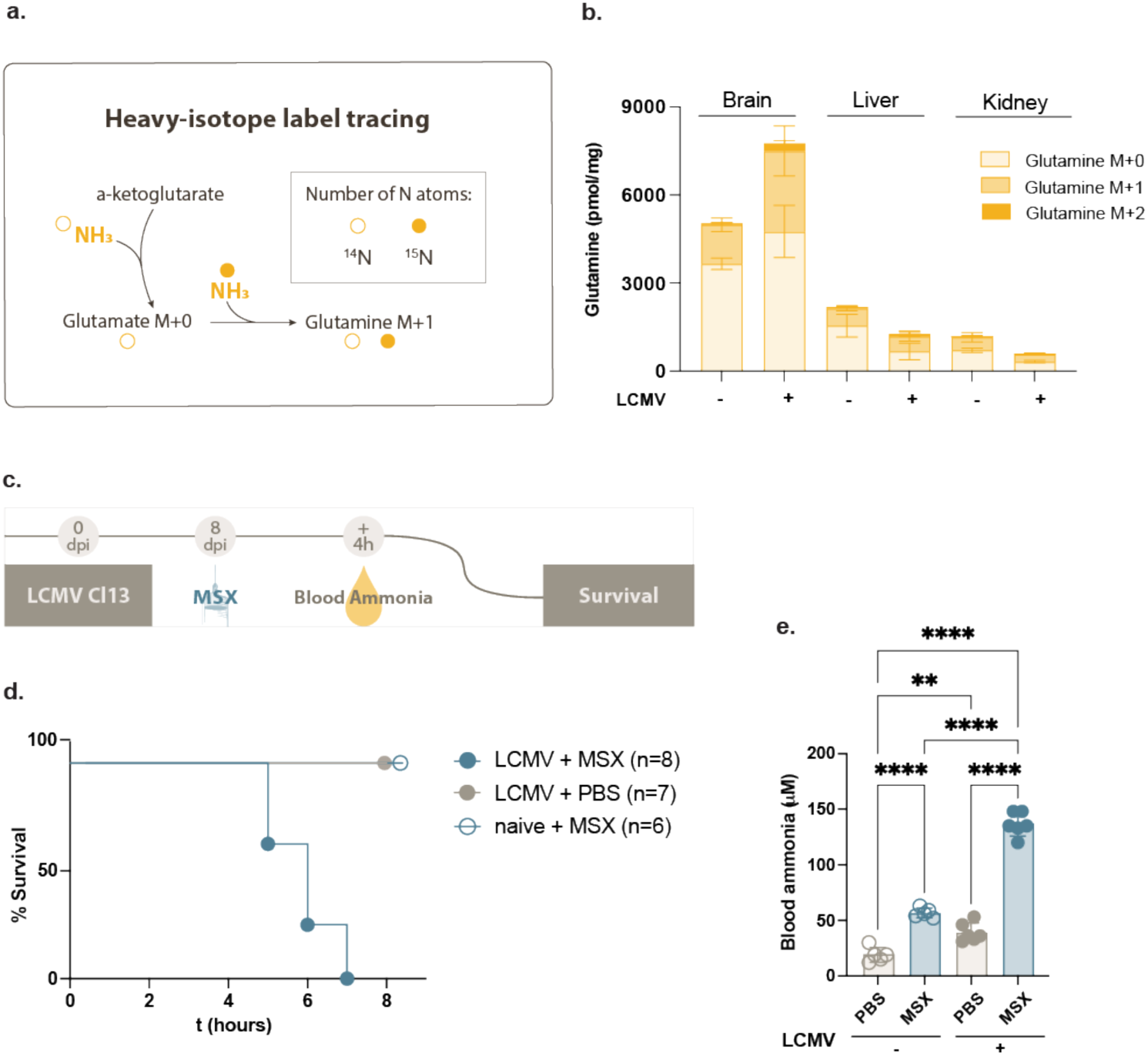
Ammonia is distributed to the brain during viral infection. (a) Schematics of heavy-isotope metabolite tracing following injection of ^15^NH_4_Cl. (b) LC-MS quantification of heavy isotope labelled glutamine in different tissues 20 min after intraperitoneal injection of ^15^NH_4_Cl at 8 days of LCMV Cl13 infection. Data represents mean±SD, n=3 mice per group, representative of two independent experiments. (c) Schematics of L-methionine-DL-sulfoximine (MSX) treatment during LCMV Cl13 infection. (d) Survival of naïve and LCMV Cl13-infected mice after injection of 50mg/kg MSX at 8 dpi. Data represent n=6-8 mice per group, pooled from two independent experiments. (e) Blood ammonia measurement at 8 days post LCMV Cl13 infection in mice pre-treated with 50 mg/kg MSX 4 h before the measurement. Data shows mean±SD, n=6 mice per group pooled from two independent experiments. Statistical analysis: one-way ANOVA with Tukey multiple comparisons test.

To perturb the conversion of ammonia into glutamine, we infected mice with LCMV Cl13 and treated them 8 days later with the glutamine synthetase (GS) inhibitor L-methionine-DL-sulfoximine (MSX). While non-infected mice tolerated the treatment without overt effects, LCMV-infected mice developed severe neurological symptoms, including seizures and impaired locomotor activity, and reached the humane endpoint between 5 and 7 h post-treatment (Figure 3c). This was accompanied by a dramatic increase in blood ammonia in infected, MSX-treated mice, compared to the non-infected, MSX-treated counterparts (Figure 3d). This data demonstrates that the brain is particularly susceptible to an increase in circulating ammonia during infection and highlights the importance of ammonia detoxification mechanisms to safeguard the brain from the toxic effects of ammonia. Additionally, it suggests a role of ammonia in modulating neuronal function during infection, similarly to what has been described in other disease models^22,44^.

### Ammonia activates the paraventricular hypothalamus and triggers a systemic stress response

Despite the well-known detrimental effects of high doses of ammonia, for example in liver disease patients^21,44^, its mechanisms of action beyond neurotoxicity remain poorly understood. To investigate the behavioral effects of acute ammonia exposure *in vivo*, we subjected mice to an open field test after an intraperitoneal injection of ammonium chloride (NH_4_Cl). We observed a rapid response characterized by a significant reduction in locomotor activity between 5 and 20 minutes after ammonia injection, and a complete recovery after 2 hours (Figure 4a). To specifically test for the effects of ammonia in neurons, we cultured primary mouse embryo cortical neurons in the presence of NH_4_Cl and performed RNA-seq on cells harvested at two different timepoints. Differential gene expression analysis revealed an upregulation of several immediate early response genes, such as *Fos*, *Jun*, *Arc*, and the *Egr* and *Nr4a* families after 2h of treatment (Figure 4b, Supplementary figure 5a). This wave of transcriptional changes wanes rapidly and is undetected at 6h (Supplementary figure 5a-b), indicating an acute neuronal response upon ammonia exposure.

**Figure 4:**
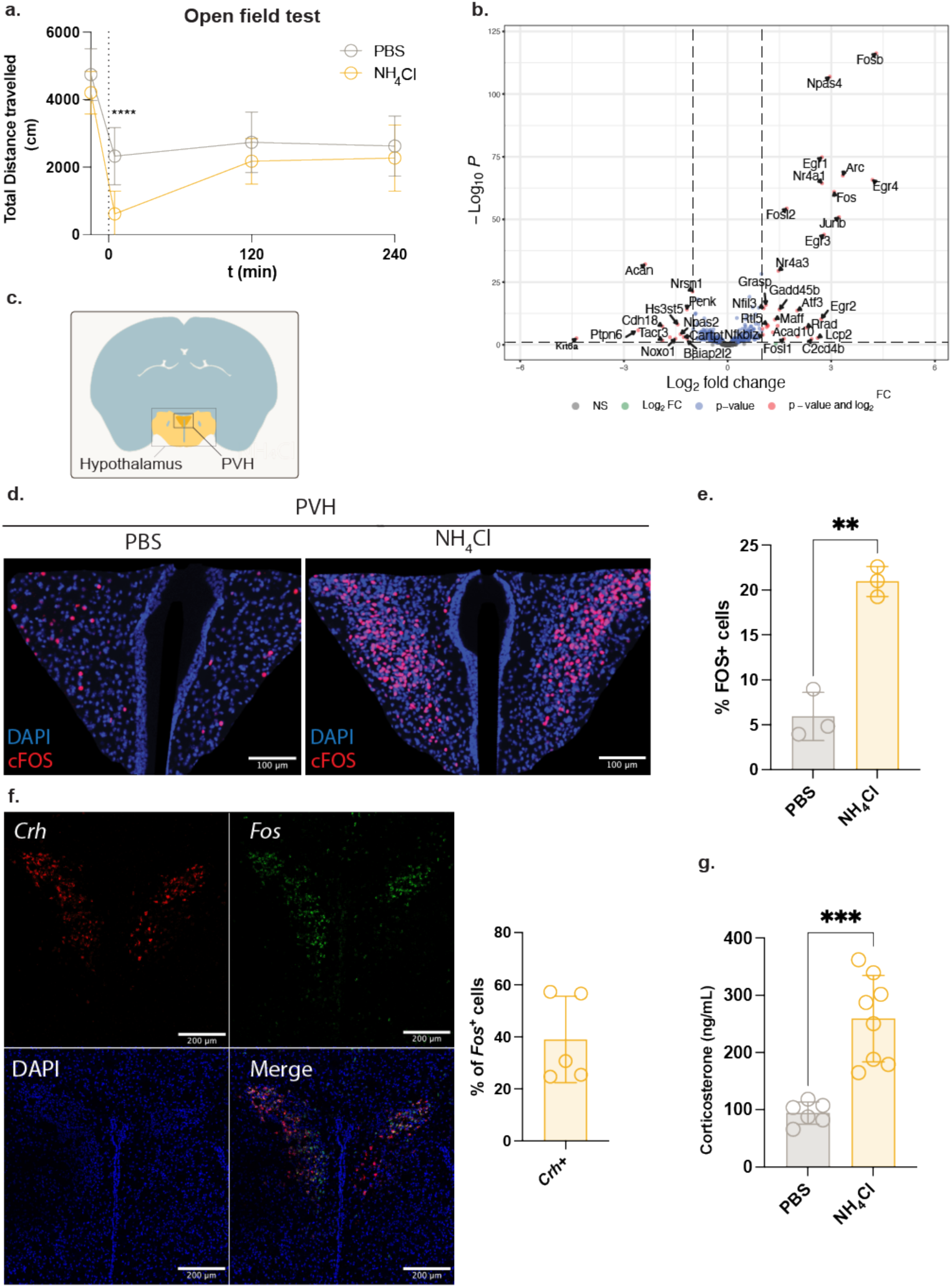
Ammonia triggers an acute behavioral response and activates the paraventricular hypothalamus (PVH). (a) Total distance travelled by mice during an open field test following an intraperitoneal injection of 200 mg/kg NH_4_Cl or PBS. For each time point mouse movement was recorded for 15 minutes. Data represents mean±SD, n=13 mice per group, pooled from two independent experiments. The dashed line (t=0) indicates the time of injection. Statistical analysis: two-way ANOVA with Šídák’s multiple comparisons test. (b) Volcano plot showing the transcriptional response of mouse embryo cortical neurons incubated with 1 mM NH_4_Cl or PBS for 2 h. n=4 mice per group. (c) Schematic of a mouse brain coronal section highlighting the hypothalamus and paraventricular hypothalamus (PVH). (d) Representative cFos immunofluorescence images of the PVH 1.5 h after an intraperitoneal injection of 200 mg/kg NH_4_Cl or PBS. (e) Quantification of cFos^+^ cells from (d). Data shows mean±SD and each dot represents the average quantification per mouse, n=3 mice per group, at least two sections per mouse, representative of two independent experiments. Statistical analysis: t-test. (f) Left: representative fluorescence *in situ* hybridization (FISH) images showing *Crh* and *Fos* staining in the PVH, 30 min after an intraperitoneal injection of 200 mg/kg NH_4_Cl. Right: quantification of *Fos+* neurons co-expressing *Crh*. Each dot represents the average quantification per mouse, n=5 mice pooled from two independent experiments. (g) Serum corticosterone measurements 30 minutes after an intraperitoneal injection of 200 mg/kg NH_4_Cl or PBS. Data shows mean±SD, n=6-8 mice per group, pooled from two independent experiments. Statistical analysis: unpaired t-test.

To determine the spatial representation of neuronal responses in the brain upon ammonia injection, we interrogated the distribution of cFos, as proxy for neuronal activation, by immunofluorescence in brain sections. We focused our analysis on the hypothalamus, a key hub for integrating peripheral immune signals and orchestrating the cardinal features of sickness behavior, including fever, anorexia, and sleep dysregulation^8,45^. We detected strong cFos signal specifically in the PVH of ammonia-treated mice but not in vehicle-treated controls (Figure 4c-e). Interestingly, this response was restricted to the PVH, as no other hypothalamic or extra-hypothalamic area showed an increase in cFos^+^ cells upon ammonia treatment (Supplementary figure 5d). Most cFos^+^ cells in the PVH of NH_4_Cl-treated mice co-localized with the neuronal marker NeuN, suggesting a neuron-specific response to ammonia (Supplementary figure 5c).

The PVH integrates inputs from the gut-brain axis to modulate downstream circuits governing neuroendocrine stress responses and behavioral adaptation^46,47^. One such pathway involves the activation of corticotropin-releasing hormone (CRH) neurons in the PVH, which stimulate adrenocorticotropic hormone (ACTH) production in the pituitary, eventually leading to the secretion of glucocorticoids from the adrenal glands into circulation. Indeed, fluorescent *in situ* hybridization (FISH) in NH_4_Cl-treated PVH sections showed a high degree of co-localization between *Fos^+^* and *Crh^+^* neurons (Figure 4f). To test whether ammonia activates a PVH-mediated systemic stress responses, we measured serum corticosterone after NH_4_Cl administration. Ammonia-treated mice showed significantly higher serum corticosterone after 30 minutes compared to vehicle-treated controls (Figure 4g), thus confirming that ammonia triggers an acute and systemic stress response.

### Pharmacological inhibition of ammonia detoxification amplifies infection-induced stress responses

Glucocorticoids play a key role in regulating immune, metabolic and behavioral responses during infection^24,48,49^. LCMV Cl13 infection leads to a CD8^+^ T cell-dependent increase in circulating corticosterone at 8 dpi (Figure 5a). Likewise, the sterile inflammation model leads to increased corticosterone levels 3 days after anti-CD3 injection (Supplementary figure 6a). To check whether elevated ammonia is associated with corticosterone production during infection, we resorted again to the MSX drug treatment to further increase blood ammonia at 8 days of LCMV Cl13 infection (Figure 3e). We measured serum corticosterone 4 hours after MSX injection and observed a two-fold increase in infected, MSX-treated mice compared to vehicle treated controls (Figure 5b). The comparison of blood ammonia and serum corticosterone levels across all experimental groups (with and without infection and MSX treatment) revealed a high degree of correlation (Pearson r = 0.8315, p < 0.0001) (Supplementary figure 6b). We next analyzed expression levels of a subset of genes, whose expression is upregulated downstream of glucocorticoid receptor (GR) activation^50,51^, in the liver and spleen. We observed a general upregulation of these genes in both analyzed organs in infected, MSX-treated mice, when compared to vehicle treated controls (Figure 5c), thus confirming a systemic GR-dependent stress response.

**Figure 5:**
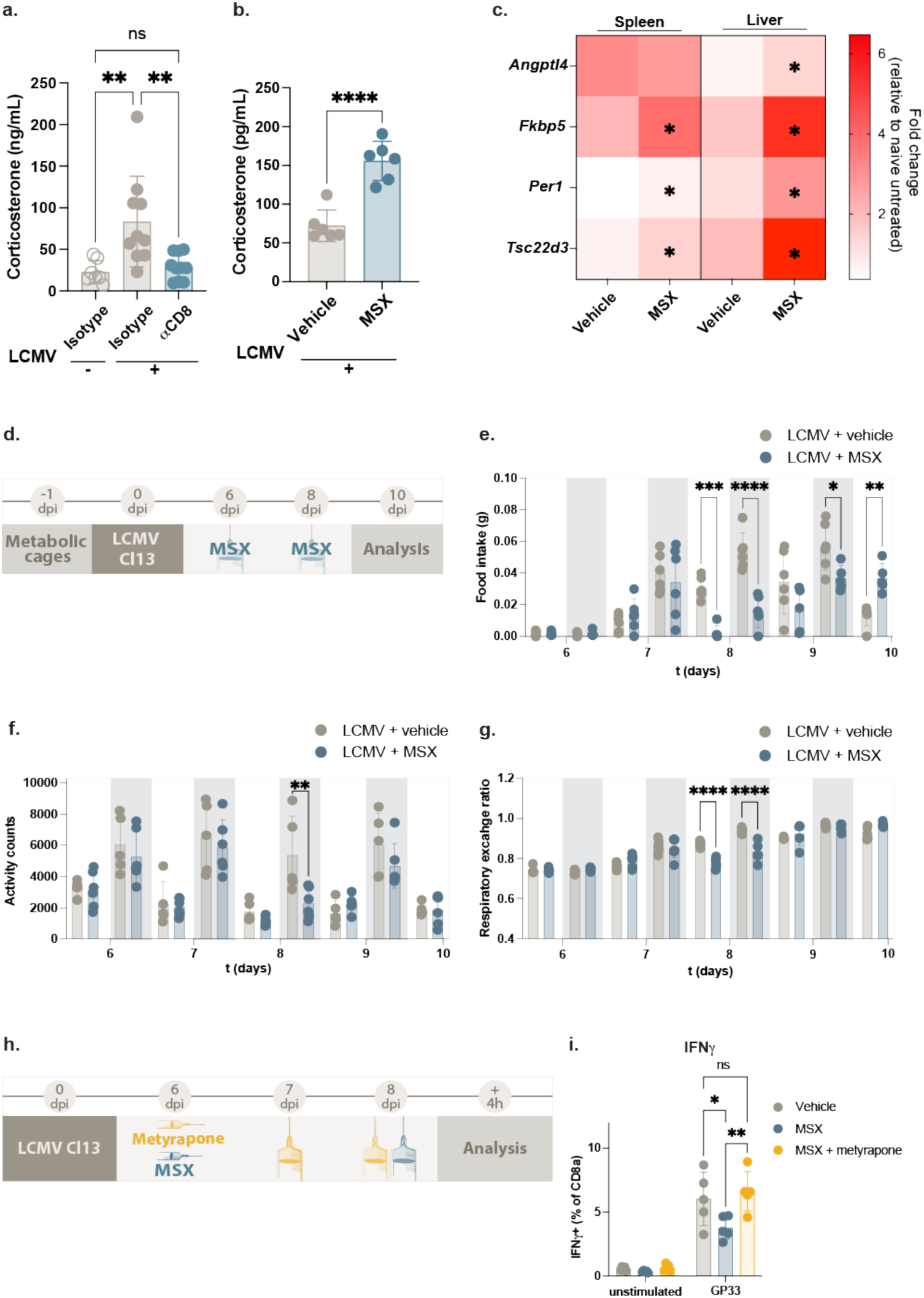
Pharmacological inhibition of ammonia detoxification amplifies infection-induced stress responses. (a) Serum corticosterone measurements at 8 days post-LCMV Cl13 infection in C57BL/6J pre-treated with an anti-CD8a antibody or an isotype control. Data shows mean±SD, pooled from two independent experiments n=7-10 mice per group. Statistical analysis: one-way ANOVA with Tukey multiple comparisons test. (b) Serum corticosterone measurements 4 hours after treatment with 50 mg/kg MSX at 8 days post-LCMV Cl13 infection. Data shows mean±SD, n=6 mice per group pooled from two independent experiments. Statistical analysis: unpaired t-test. (c) Expression analysis of glucocorticoid receptor-dependent genes obtained by quantitative PCR in spleen and liver 4 hours after treatment with 25 mg/kg MSX at 8 days post-LCMV Cl13 infection. The heat map displays the fold change expression relative to non-infected, vehicle-treated mice, n= 4-5 mice per group. Statistical analysis: multiple unpaired t-tests with Holm-Šídák correction. (d) Schematic summary of sublethal MSX administration in metabolic cages. (e-g) Metabolic cage recordings in C57BL/6J mice infected with LCMV Cl13 and treated with 25 mg/kg MSX: (e) food intake, (f) activity counts, and (g) respiratory exchange ratio (RER). Data shows the average recordings over 15 minute-periods during the active (dark) and rest (light) phases. Shaded areas indicate the dark cycle. n=5-6 mice per group, pooled from two independent experiments. (h) Schematic summary of MSX and metyrapone treatments during LCMV Cl13 infection. (i) Percentage of IFNγ producing T cells in unstimulated or GP33-stimulated splenocytes. Mice had been previously infected with LCMV Cl13 and treated with MSX and/or metyrapone. Data shows mean±SD, n=9-10 mice per group pooled from two independent experiments. Statistical analysis: two-way ANOVA with Tukey multiple comparisons test.

Considering the far-reaching effects of glucocorticoids during infection, we hypothesized that a sublethal MSX treatment would have an impact on the host response to LCMV Cl13. To obtain an overview of the metabolic and behavioral effects of MSX treatment, we placed mice in metabolic cages with continuous monitoring over the infection and MSX treatment period (Figure 5d). LCMV Cl13 infection in vehicle-treated mice led to the expected sickness behavior starting at 6 dpi, which results in decreased locomotion and food intake, followed by a recovery phase starting on the night of day 7 (Figure 5e-f, Supplementary figure 6c-d). In contrast, MSX treatment caused a delay in recovery with reduced food intake and activity until 9 dpi (Figure 5e-f). This was accompanied by prolonged metabolic reprogramming upon MSX treatment, as indicated by a decrease in respiratory exchange ratio (RER) that persisted until 9 dpi (Figure 5g). MSX treatment in non-infected mice led to a slight decrease in food intake (Supplementary figure 6c), along with a transient decrease in activity and RER (Supplementary figure 6 d-e). This data shows that pharmacological inhibition of ammonia detoxification leads to behavioral changes, which, in the context of viral infection, result in a delayed recovery from sickness behavior.

We next explored whether ammonia accumulation during infection enhances the immunosuppressive effect of glucocorticoids. To this end, mice infected with LCMV Cl13 were treated with MSX and the glucocorticoid synthesis inhibitor metyrapone, and T cell function was analyzed 4 hours after the last drug administration (Figure 5g). Analysis of splenic T cells after stimulation with the viral peptide GP33 showed a significant reduction in IFNγ production upon MSX treatment, which was reverted when mice were co-treated with MSX and metyrapone (Figure 5h, Supplementary figure 6e). Together, our data shows that ammonia accumulation during infection contributes to the activation of CRH^+^ neurons in the PVH and increased corticosterone release, which plays a critical role in the behavioral, metabolic and immune response to infection.

## Discussion

In this study, we uncover ammonia as systemic signal driving the interplay between T cells, the gut microbiota and the brain during infection, leading to activation of the hypothalamic stress response. Under homeostatic conditions, both the gut microbiota and mammalian cells produce low levels of ammonia, which is then detoxified in the liver urea cycle and in extra-hepatic tissues via glutamine synthesis^20^. We propose that during chronic viral infection with LCMV Cl13 in mice, T cell-mediated inflammation induces gut dysbiosis leading to an increase in ammonia production primarily driven by shifts towards higher abundances and activity of proteolytic clostridial taxa. Considering the transient anorectic behavior of LCMV-infected mice, this data suggests that host protein-derived nitrogen utilization by such bacterial taxa is a possible source of ammonia release. This increase in ammonia is further aggravated by the disruption of the hepatic urea cycle in infected mice^23^. Ammonia is then sensed by the PVH and results in activation of the hypothalamus-pituitary-adrenal (HPA) axis and an increase in circulating corticosterone. Our results reveal a signaling effect of ammonia, contributing to a systemic stress response, while simultaneously exposing the vulnerability of infected mice to the toxic effects of ammonia in the brain. Treatment with a high dose of the glutamine synthetase inhibitor MSX, which prevents extra-hepatic ammonia detoxification^52,53^, causes severe neurological symptoms in infected mice, coinciding with high ammonia levels that would otherwise be safely eliminated in non-infected mice. CD8^+^ T cells hold important roles beyond classical antiviral function and can have a dramatic impact in physiology by rewiring parenchymal cell metabolism and causing immunopathology in tissues like the liver and the gut ^11,23,24,32,54^. Our results demonstrate that T cells are necessary and sufficient to increase blood ammonia even in the absence of viral infection. We propose that T cell infiltration in the intestine and consequent dysbiosis are main drivers of hyperammonemia. In support of this notion, depletion of neomycin-sensitive gut microbiota in LCMV Cl13 prevents hyperammonemia but does not prevent liver immunopathology and urea cycle dysfunction. In parallel, we observed elevated blood ammonia 72 hours, but not 24 hours, after anti-CD3 injection. The anti-CD3 sterile inflammation model is characterized by cytokine storm-like systemic inflammation within hours of treatment, followed by gut-specific T cell infiltration and intestinal immunopathology over the following days^27,28,55,56^. The delayed increase in blood ammonia therefore coincides with the onset of tissue-specific pathology. Together, these results suggest a central role of T cell-mediated gut pathology in hyperammonemia. The effects of ammonia are highly dose- and context-dependent, due to its chemical properties that allow for rapid transport, incorporation in secondary metabolites, and elimination. At high doses, ammonia has neurotoxic effect^44^, posing a life-threatening risk for patients with compromised liver function^57^. At moderate concentrations, ammonia acts through multiple signaling mechanisms reminiscent of other small, gaseous molecules such as nitric oxide and carbon monoxide^16^. These mechanisms, which have only recently begun to be explored, include a neuroprotective effect against depression^22^, multiple effects on T cells^58,59^ with important implications in cancer progression^60,61^, and an indirect gut-brain communication axis to control feeding behavior^62^.

Our study uncovers a previously unrecognized role of ammonia in activating hypothalamic stress responses and increasing circulating levels of glucocorticoids. This observation matches anecdotal evidence from farm animals, which are often exposed to high ammonia levels due to overcrowded housing. In this specific context, chronic exposure to ammonia has been associated with increased levels of stress hormones, such as CRH, as well as behavioral changes^63–65^.

In humans, hyperammonemia is primarily associated with liver disease and cases of extra-hepatic origin are rare and poorly understood^66^. In the infection context, the recent COVID-19 pandemic revealed several cases of elevated blood ammonia without known liver disease^67–69^. This is likely an underdiagnosed phenomenon, as blood ammonia is not routinely measured in patients without prior disease. A broader characterization of ammonia levels in infected patients and its association with circulating glucocorticoids might help shed light into this overlooked aspect of disease. During infection, an acute increase in endogenous glucocorticoids provides an essential immunosuppressive effect that helps balancing the immune response and avoiding immunopathology^70,71^. Furthermore, post-acute infection syndromes, such as long COVID, have been associated with persistently low levels of cortisol, which is thought to contribute to disease severity^72–74^. Even though inflammatory mediators such as cytokines have been proposed as the main drivers of infection-induced stress responses^7,75^, gut microbiota also play a key role in stress regulation^76^. Specifically, the PVH senses microbial stimuli and initiates the signaling cascades that result in glucocorticoid synthesis^47^.

The downstream effects of ammonia-induced PVH activation have potentially far-reaching consequences during and after infection, warranting further investigation in future studies. For example, PVH activation and the consequent increase in corticosterone have a negative effect on social activity in mice^46^, which may explain social isolation as part of the sickness behavior in the context of infection. This notion is supported by our observation that sublethal MSX treatment during LCMV Cl13 infection prolongs sickness behavior and delays recovery from anorexia. Additionally, we found that MSX treatment suppresses antiviral T cell function in in a glucocorticoid-dependent manner, suggesting that ammonia indirectly restrains T cell activity, therefore preventing excessive immunopathology. In light of these observations, our work reveals a new role of gut-derived ammonia in boosting the systemic adaptation to infection via PVH stimulation, while opening new avenues for the use of microbiome-targeted therapies to alleviate dysregulated stress responses.

## Acknowledgements

The authors would like to thank Maria Rescigno (CeMM) and Michael Wagner (University of Vienna) for scientific discussion. This work was supported by the European Research Council under the European Union’s Seventh Framework Program and Horizon 2020 research and innovation program, grant agreements no. 677006 (“CMIL”) to AB and no. 101028971 (“AmmoniaVir”) to HGC. AB received grants from the Austrian Science Fund (FWF) no. 35806, no. 36646 and Cluster of Excellence Microbiomes drive Planetary Health (10.55776/COE7). RNA sequencing was performed by the Biomedical Sequencing Facility at CeMM Research Center for Molecular Medicine of the Austrian Academy of Sciences, while 16S rRNA gene amplification and sequencing was performed by the Joint Microbiome Facility of the Medical University of Vienna and the University of Vienna. We thank the NIH Tetramer Core Facility (NIH Contract 75N93020D00005 and RRID:SCR_026557) for providing tetramer reagents.

## Methods

### Mice

All animal experiments were approved by the Austrian Federal Ministry Women, Science and Research under licenses BMWFW-66.009/0361-WF/V/3b/2017, 2020-0.406.011, and 2024-0.363.541.

Mice were kept and bred at the core facility for laboratory animal breeding and husbandry of the Medical University of Vienna, under specific pathogen-free (SPF) conditions and in individually ventilated cages. Wild-type C57BL/6J mice were bred in-house or purchased from Janvier laboratories. *Ifnar1*^−/−^ (JAX 032045)^77^ and *Rag2^−/−^* (JAX 008449)^78^ mice were bred in-house and kept on C57BL/6J background. All mice were used between 8 and 15 weeks of age. Within each experiment mice were age and sex matched.

### Infection and inflammation models

For LCMV Cl13 infection, mice received an intravenous injection of 2×10^6^ focus forming units (ffu) via lateral tail vein. Viral stocks were generated in-house using BHK-21 cells as previously described^79^. Control mice were injected with PBS.

For the anti-CD3 sterile inflammation model, model mice received an intraperitoneal injection of 50 μg of anti-mouse CD3ε clone 145-2C11 (BioXcell catalog #BE0001) diluted in PBS. Control mice received the same amount of polyclonal Armenian hamster IgG isotype control (BioXcell catalog #BE0091).

For poly(I:C) treatment, mice received an intraperitoneal injection of 10 mg/kg poly(I:C) high molecular weight (Invivogen catalog #tlrl-pic). Poly(I:C) stocks were prepared by dissolving in saline and incubating at 65 °C for 10 min. Control mice received a PBS injection.

Mice were sacrificed by cervical dislocation at the indicated time points, tissues were immediately snap frozen in liquid nitrogen and stored at −80 °C.

### Acute ammonia treatment

Mice were habituated to handling stress by repeated manual restraining and injection of 100 μL PBS daily for at least three days prior to the experiment. NH_4_Cl (Sigma catalog #A4514) was dissolved in PBS, sterilized by filtration with a 0.2 μm syringe filter and injected intraperitoneally at 200 mg/kg. Control mice received an intraperitoneal injection of PBS. Mice were sacrificed by an intraperitoneal injection of 300 mg/kg ketamine and 20 mg/kg xylazine. The timepoints of brain harvest were selected to reflect the different temporal dynamics of RNA and protein expression: 30 minutes for RNA analysis by FISH and at 90 minutes for protein analysis by immunofluorescence.

### Pair feeding

Mice were co-housed and the average food intake of infected or anti-CD3 treated mice was recorded daily. For pair feeding, naïve mice received the same amount of food consumed by infected animals on a per day basis.

### Drug treatments

For gut microbiota depletion, mice were treated with 1 mg/mL ampicillin (Roth catalog #K029.1), 0.5 mg/mL metronidazole (Santa Cruz catalog #sc-204805), 1 mg/mL neomycin (Santa Cruz catalog #sc-3573A), 0.25 mg/mL vancomycin (Sigma catalog #SBR00001), 1 mg/mL gentamycin (Santa Cruz catalog #sc-203334A), alone or in combination, in the drinking water. Water was supplemented with 4 mg/mL sucralose (Sigma catalog #69293) to mask the bitter taste of the antibiotics. Antibiotics and sucralose were dissolved in tap water and sterilized by filtration with a 0.2 μm filter. Treatment was started 10 days before infection and maintained throughout the experiment. Antibiotics were prepared fresh and replaced twice a week. In addition to the treatment in the drinking water, mice received at the start of the experiment and 12 to 16 h before harvest an oral gavage of 200 μL containing 5 mg/mL ampicillin, 2.5 mg/mL metronidazole, 5 mg/mL neomycin, 1.25 mg/mL vancomycin, and 5 mg/mL gentamycin dissolved in PBS.

The glutamine synthetase inhibitor methionine-DL-sulfoximine (MSX, MedChem Express catalog # HY-B1692) was dissolved in PBS and injected intraperitoneally at 8 days post LCMV Cl13 infection at a dose of 50 mg/kg (lethal dose) or 25 mg/kg (sublethal dose). Control mice received a vehicle (PBS) injection. After injection, mice were monitored every hour and scored for signs of neurological problems such as severe apathy, disrupted balance, or spasms. Animals were immediately euthanized upon reaching the humane end point established in the approved ethics protocols.

The corticosterone synthesis inhibitor metyrapone (Tocris catalog # 3292) was dissolved in 0.5% carboxymethylcellulose (CMC) and injected intraperitoneally daily between 6 and 8 dpi at a dose of 50 mg/kg. Control mice received a CMC injection. For metyrapone and MSX co-treatments, metyrapone was injected 30 minutes before MSX.

The urease inhibitor acetohydroxamic acid (AHA, Sigma catalog # 159034) was dissolved in PBS and administered via oral gavage daily between 4 dpi and 7 dpi at a dose of 250 mg/kg. Control mice were gavaged with PBS. The urease inhibitor flurofamide (FFA, RCD systems catalog #0478) was dissolved in DMSO and diluted in PBS for a final DMSO concentration of 7.5%. The resulting solution was injected intraperitoneally at 7 days post LCMV Cl13 infection at a dose of 6 mg/kg. Control mice received an injection of 7.5% DMSO in PBS.

### Blocking and depletion antibodies

For CD8^+^ T cell depletion, mice received an intraperitoneal injection of 200 μg anti-mouse CD8a clone YTS169.4 (BioXcell catalog #BE0117) two and one day before infection with LCMV Cl13. Control mice received a rat IgG2b isotype control clone LTF-2 (BioXcell catalog #BE0090). For CXCR3 blocking, mice received an intraperitoneal injection of 200 μg anti-mouse CXCR3 clone CXCR3-173 (BioXcell catalog #BE0249) at days −1, 2, and 5 post-LCMV Cl13 infection. Control mice received a polyclonal Armenian hamster IgG isotype control (BioXcell catalog BE0091).

### Metabolic cages

Mice were housed individually in PhenoMaster cages (TSE systems). Mice were transferred to the metabolic cages 24 h before the start of the experiment to allow for acclimatization and obtain baseline measurements. Activity counts, food intake, oxygen consumption and exhaled CO_2_ were recorded in 10 to 15-minute cycles.

### Open field test

Locomotor and exploratory behavior were evaluated in an open field test divided in five trials of 15 min each. The first trial (baseline) was performed 1 h before NH_4_Cl treatment, the second 5 min after injection and the subsequent trials in two-hour intervals. Mice were placed in a 27.5 cm × 27.5 cm arena and allowed to freely explore the environment for each trial. Movement was recorded and quantified using integrated laser beams (Activity Monitor v5, MedAssociates, Fairfax, VT, USA).

### Biochemical measurements

Ammonia was measured from freshly collected tail vein blood using a PocketChem™ BA PA-4140 Blood Ammonia Meter (Arkray) according to the manufacturer’s instructions. Briefly, 20 μL whole blood was collected from the tail vein, added to an ammonia test strip, incubated for 3 minutes at room temperature and immediately measured.

For biochemical measurements in serum, blood was obtained by cardiac puncture and allowed to clot at room temperature, followed by centrifugation at 10,000 x g for 5 minutes at 4 °C. Serum was collected and stored at −80 °C until further analysis. Alanine aminotransferase (ALT) was measured using a Cobas C311 analyzer (Roche). Corticosterone was determined in serum collected during the light phase (Zeitgeber time, ZT 1 to 5) using an ELISA Kit (Enzo Life Sciences catalog #ADI-901-097) according to the manufacturer’s instructions and using a 1:40 serum dilution.

Urease activity was determined in freshly collected fecal pellets using a colorimetric Urease Activity Assay Kit (Abcam catalog #ab204697). Briefly, one fecal pellet was weighed and homogenized in 1 mL PBS with a 5-mm stainless-steel bead (Qiagen catalog #69989) and 0.1mm glass beads (Cell Disruption Media, Scientific Industries, Inc catalog #888-850-6208) using a TissueLyser II (Qiagen) at 30 s^−1^ for 2×30 s. Homogenates were centrifuged at 14,000 x *g* for 10 min at 4 °C. Supernatants were diluted 1:2 with assay buffer and then assayed according to the manufacturer’s protocol. Absorbance was measured against a NH_4_Cl standard curve (0-20 nmol) at 670 nm and measurements normalized to fecal weight.

### Flow cytometry

Spleens were collected in PBS and kept on ice until further processing. Upon mechanical dissociation though a 40 μm cell strainer, cells were resuspended in 5 mL PBS, centrifuged at 400 x *g* for 5min at 4°C, and resuspended in PBS for staining.

Intestinal T cells were prepared as described^80^. Briefly, small intestine (including the terminal portion of the ileum) and large intestine (including both the cecum and colon) were cleaned of fat, lymphoid structures and intestinal contents and incubated for 20 min at 37 °C, 250 rpm in a buffer containing PBS with 2% fetal bovine serum (FBS), 10 mM HEPES, 1 % penicillin/streptomycin, 1 % L-glutamine, 1 mM EDTA and 1 mM dithiothreitol to remove intraepithelial lymphocytes. The remaining intestinal tissue was rinsed in PBS and further digested for 20 min at 37 °C, 250 rpm in 50-mL tubes containing 3x one-quarter-inch ceramic beads (MP Biomedicals catalog #15582384) and 25 mL collagenase solution (RPMI 1640 with 2 % FBS, 10 mM HEPES buffer, 1 % penicillin/streptomycin, 1 % L-glutamine, 1 mg.ml^−1^ collagenase D (Roche catalog catalog #11088858001) and 1 U.ml^−1^ DNase I (Sigma catalog #11284932001)). The resulting lamina propria lymphocytes (LPL) were passed through a 100-μm strainer and centrifuged at 400 *g* for 5 min at 4 °C. Cell suspensions were then washed by centrifugation in 40 % Percoll (Cytiva catalog #17-0891-02) in PBS. Cell pellets were resuspended in FACS buffer (PBS with 2% FBS and 2 mM EDTA) for staining.

For surface staining, cells were incubated with CD16/32 FcR-Block, Zombie UV™ Fixable Viability dye, LCMV virus specific tetramers (PE-GP33, APC-Np396 and FITC-Gp276 obtained from the NIH Tetramer Core Facility, US), and surface antibodies (CD8a-Alexa Fluor 700 or BV711, TCRb-BUV395, CD45-PerCP, CD103-Pacific Blue, CD44-Alexa Fluor 700, CXCR3-BV421) diluted in FACS buffer, for 20 minutes at room temperature. Cells were subsequently washed and fixed with 4% paraformaldeyhde (PFA) for 10 min at room temperature. For intracellular staining, cells were incubated with 1 μg/mL GP33 peptide (Anaspec catalog #AS-61669) for 4 h at 37 °C in RPMI medium supplemented with 10 % FBS, 1 % Penicillin/Streptomycin, 50 μM β-mercaptoethanol and Protein Transport Inhibitor Cocktail (eBioscience catalog# 00-4980-03). Cells were subsequently washed in FACS buffer, stained for surface markers, and fixed as described above. Following permeabilization with the eBioscience™ Foxp3/ Transcription Factor Staining Set (Invitrogen catalog #00-5523-00), the following intracellular proteins were stained for 90 min at 4 °C: IFNγ-Pe-Cy7 or RY586 and TNF-APC. Flow cytometry data was acquired in a Cytek Aurora spectral flow cytometer and analyzed in FlowJo 10.

### RNA extraction and quantitative PCR

Tissues were homogenized in Qiazol using a TissueLyser II (QIAGEN). RNA was extracted using QIAzol (Qiagen catalog #79306) and chloroform (Carl Roth catalog #3313.1) following manufacturer’s instructions. The resulting RNA was quantified using a Nanodrop and 500 ng RNA were used for cDNA synthesis using random primers and the LunaScript® RT SuperMix (New England Biolabs catalog #M3010X) according to the manufacturer’s instructions. Real time PCR was performed with the Luna Universal Probe qPCR Master Mix (New England Biolabs catalog #M3004E) and the following Taqman Gene expression Assays (Thermo Fischer Scientific, catalog #4331182): *Gls* (Mm01257297_m1), *Glul* (Mm00725701_s1), *Glud1* (Mm00492353_m1), *Otc* (Mm00493267_m1), *Angptl4* (Mm00480431_m1), *Fkbp5* (Mm00487406_m1), *Per1* (Mm00501813_m1), *Tsc22d3* (Mm00726417_s1), and the housekeeping gene *Eef1a*1, (Mm01973893_g1).

### Metabolomics

For heavy isotope tracing, mice received an intraperitoneal injection of 200 mg/kg ^15^N-NH_4_Cl (Sigma catalog #299251) at 8 dpi. Mice were sacrificed by cervical dislocation 20 minutes after injection and tissues were frozen in liquid nitrogen and stored at −80 °C until further analysis.

For cecal metabolomics, mice were sacrificed by cervical dislocation at 8 dpi, the cecal contents collected aseptically and frozen in liquid nitrogen.

Tissue samples were weighed, 3 µL MS-grade methanol were added per mg tissue and samples were homogenized using a Precellys 24 tissue homogenizer with a Precellys CK14 lysing kit (Bertin). The homogenization was performed at 0-4 °C at 600 rpm for 30 sec, with a rest time of 5 sec, repeating the cycle two times. Tubes were then centrifuged at 10,000 x g for 10 min at 4 °C. Fifty microliters of supernatant were collected, transferred to a 96-well hydrophobic filter plate (MultiScreen Solvinert, hydrophobic PTFE, 0.45 µm, MSRPN0410, Milipore) and mixed with 20 µL of an isotopically labelled internal standard mixture. Then, 270 µL of methanol were added and the plate was shaken for 20 min at 450 rpm. Afterwards, the sample extract was collected in an attached 96-well plate by centrifuging the filter plate for 10 min at 1000 g. Sample extracts were used for LC-MS analysis.

A Vanquish UHPLC system (Thermo Scientific) coupled with an Orbitrap Q Exactive (Thermo Scientific) mass spectrometer was used for LC-MS analysis. The chromatographic separation was carried out on an ACQUITY UPLC BEH Amide, 1.7 µm, 2.1×100 mm analytical column (Waters) equipped with a VanGuard: BEH C18, 2.1×5 mm pre-column (Waters). The column was maintained at 40 °C and the sample injection volume was 2 µL. The mobile phase consisted of phase A - 0.15 % formic acid (v/v) and 10 mM ammonium formate in water and phase B - 0.15 % formic acid (v/v) in 85 % acetonitrile (v/v) and 10 mM ammonium formate. The gradient elution with a flow rate of 0.4 mL/min was performed for a total analysis time of 17 min. The Orbitrap Q Exactive (Thermo Scientific) mass spectrometer was operated in a positive electrospray ionization mode, spray voltage 3.5 kV, auxiliary gas heater temperature 400 °C, capillary temperature 350 °C, auxiliary gas flow rate 12, sheath gas flow rate 50. The metabolites of interest were analyzed using a full MS scan mode, scan range m/z 50 to 400, resolution 35000, AGC target 1e6, maximum IT 50 ms. The Skyline software (64-bit, vers. 25.1.0.237) was used for the data processing. A ten-point linear calibration curve with internal standardization and 1/x2 weighing was constructed for metabolite quantification.

For statistical analysis, only metabolites detected in more than 4 samples and at least 3 replicates of each condition were considered. Missing values were replaced by 1/5 of the limit of detection of each metabolite; detected, but not quantified, values, by the limit of detection. Metabolites for which no quantification by an internal standard was performed, were divided by the input cecum weight. A Welch t-test (base R t.test function) on the log-transformed metabolite concentrations was used to test for significant differences between conditions. The False Detection Rate (FDR) was controlled using the Benjamini-Hochberg method.

### 16S rRNA gene amplicon sequencing and copy number determination

Cecal contents were collected at 8 dpi, immediately frozen in liquid nitrogen and kept at −80 °C until further analysis. DNA extraction, library preparation and sequencing were performed at the Joint Microbiome Facility (Vienna) under JMF Project ID JMF-2510-01 as previously described^81^.

Briefly, DNA was extracted from cecal content using the QIAamp Fast DNA Stool Mini Kit (Qiagen), followed by amplification of the bacterial and archaeal 16S rRNA gene V4 region using primers 515F and 806R^82,83^, barcoding PCR, library preparation and sequencing on Illumina MiSeq (V3 chemistry, 600 cycles). Amplicon pools were extracted from the raw sequencing data using the FASTQ workflow in BaseSpace (Illumina) with default parameters, and amplicon pools were further demultiplexed with the python package demultiplex (Laros JFJ, github.com/jfjlaros/demultiplex) allowing one mismatch for barcodes and two mismatches for linkers and primers. Amplicon sequence variants (ASVs) were inferred using the DADA2 R package v1.36^84^ applying the recommended workflow ^85^. FASTQ reads 1 and 2 were trimmed at 220 nt and 150 nt with allowed expected errors of 2 on both reads. ASV sequences were against the SILVA database SSU Ref NR 99 release 138.2^86^.

For absolute quantification of 16S rRNA gene copy numbers, digital (d)PCR on a QIAcuity Digital PCR System, using the EvaGreen (EG) PCR Kit, the same DNA extracts used for 16S rRNA gene amplicon sequencing, and the same 16S rRNA-gene V4-targeted primers (515F/806R, with 0.4 µM per reaction for each primer) was performed. Cycling conditions were as follows: 2 min initial denaturation at 95° C, followed by 40 cycles of 30 s/95°C denaturation, 30 sec/52°C annealing and 30 s/72° C elongation, and a final 5 min incubation at 40° C were performed. Fluorescence was read out after each cycle elongation step. dPCR data was converted to 16S rRNA gene copy numbers per gram cecal content and used for absolute abundance conversion of the compositional 16S rRNA gene amplicon sequencing output in all downstream analyses.

Downstream analyses were performed using R v4.5.3 and packages SummarizedExperiment v1.40, SingleCellExperiment v1.24, TreeSummarizedExperiment v2.18^87^, mia v1.18 (https://github.com/microbiome/mia), vegan v2.7-2, phyloseq v1.54^88^, microbiome v1.32 (http://microbiome.github.io), and microViz v0.12.7^89^. Alpha diversity (i.e. Chao 1 richness estimator) was calculated using R packages vegan and mia. In order to visualize the relationship between the metabolome and microbiome datasets, we generated a redundancy analysis (RDA) biplot using R package vegan. Prior to plotting, the dPCR-normalized count matrix was aggregated by genus and subsequently filtered based on prevalence, retaining only genera detected in at least two replicates per group. A subset of the most differentially abundant metabolites was selected for plotting, which included creatinine, 3-hydroxy-DL-knurenine, L-alanine, L-carnosine, choline, L-phenalalanine, L-arginine, L-threonine, L-glutamine and glutamic Acid.

To identify differentially abundant bacterial genera, we performed a differential abundance analysis using DESeq2 on the same prevalence-filtered count matrix aggregated at the genus level. Since absolute dPCR-normalized abundances are not a suitable input format for DESeq2, a relative abundance matrix was used for model fitting, and dPCR determined abundances were provided as external size factor. Differentially abundant genera were identified between groups, and p-values were adjusting for multiple testing using the false discovery rate (FDR) correction. For visualization, dPCR-normalized abundances (copies/biomass) were used to illustrate the magnitude of differences between treatments.

The 16S rRNA gene amplicon sequencing data was deposited to NCBI (https://www.ncbi.nlm.nih.gov/) under the BioProject accession number PRJNA1454696.

### Primary neurons

Cortical neurons were isolated from E18.5 mouse embryos as previously described^90^. At DIV5 (i.e., 5 days after plating), cells were incubated for 2 h or 6 h with 1mM NH_4_Cl dissolved in neurobasal medium (Thermo Fisher catalog #21103049). After incubation, the medium was aspirated and cells were immediately lysed in Qiazol and stored at −20 °C before RNA extraction, as described above.

### RNA-seq

Total RNA was quantified using the Qubit® RNA BR Assay Kit (Thermo Fisher Scientific catalog #Q10211) and the RNA integrity number (RIN) was determined using the RNA 6000 Pico Kit (Agilent catalog #5067-1513) on a 2100 Bioanalyzer High-Resolution Automated Electrophoresis instrument (Agilent catalog #G2939A/B). RNA-seq libraries were prepared with the NEBNext® Poly(A) mRNA Magnetic Isolation Module E7490, the NEBNext® Ultra™ II Directional RNA sample preparation kit E7760 and NEBNext® Multiplex Oligos for Illumina (Unique Dual Index UMI Adaptors RNA) (New England Biolabs). NGS library concentrations were quantified with the Qubit® 1X dsDNA HS Assay Kit (Thermo Fisher Scientific catalog #Q32856), and the size distribution was assessed using the High Sensitivity DNA Kit (Agilent catalog #5067-4626) on a 2100 Bioanalyzer instrument. Before sequencing, sample-specific NGS libraries were diluted and pooled in equimolar amounts. Expression profiling libraries were sequenced on a HiSeq 3000 instrument (Illumina) following a 50-base-pair, single-end recipe.

Raw read quality was checked using FastQC (v0.11.9) (https://www.bioinformatics.babraham.ac.uk/projects/fastqc). Gene expression counts were estimated from raw reads using the alignment free method implemented in Salmon (v.1.4.0)^91^ against the mm10_cdna salmon index obtained via refgenie (v0.7.0)^92^.

For identifying differentially expressed genes, read counts predicted by Salmon were imported into R via tximport (v1.38.2)^93^, and genes were annotated with biomaRt using the ensemble mmusculus_gene_ensembl data (v.2.60.0)^94^. Only genes with at least 10 reads and 1 count per million reads (cpm) across at least 3 samples were kept for further analysis. Differential expression was assessed using the nbinomWaldTest function of DESeq2 (v1.46.0)^95^ with shrinkage of fold changes using the lfcShrink function with an adaptive shrinkage estimator (ashr) and using independent hypothesis weighting (IHW) for multiple testing adjustment. Genes with a P value cutoff of 0.1 and an absolute log2 fold change greater of equal 1 were deemed significantly differentially expressed.

All statistical analysis were performed in R (v4.4.1) (R Core Team, 2024), graphics were created using ggplot2 (v4.0.1) and EnhancedVolcano (v1.22.0) for Volcano plots. The data have been deposited in the European Nucleotide Archive (ENA) at EMBL-EBI under accession number PRJEB113181 (https://www.ebi.ac.uk/ena/browser/view/PRJEB113181).

### Immunofluorescence

Mice were sacrificed with an intraperitoneal injection of 300 mg/kg ketamine and 20 mg/kg xylazine, perfused transcardially with PBS and fixed *in-situ* with 4 % PFA diluted in PBS. Brains were carefully extracted and post-fixed in 4 % PFA for 24 h at 4 °C. The fixation solution was subsequently replaced by 30 % sucrose for 48 h at 4 °C. Brains were then embedded in optimal cutting temperature (OCT) compound, frozen in dry ice and stored at −80 °C until further processing.

Cryosections were obtained in 30 µm coronal slices using a cryostat (Leica® catalog # CM1950) and stored at −20 °C in cryoprotectant solution (30 % glycerol, 30 % ethylene glycol, and 40 % PBS 1x). Slices were stained with primary antibodies for rabbit anti-phospho-c-Fos (1:1000, Cell Signalling Technology catalog #5348) and mouse anti-NeuN (1:500, Merck Millipore catalog # MAB377). For secondary antibodies, donkey anti-rabbit 594 (1:1000, Jackson ImmunoResearch Laboratories, catalog #JAC711587003) and donkey anti-mouse 647 (1:1000, Invitrogen catalog # 15980296) were used. Slide scans were obtained in a Vectra Polaris PhenoImager or Axiovert 200M Fluorescence/Live cell Imaging Microscope with 10x and 20x objectives (Carl Zeiss AG) and images were analyzed using FIJI 2.16 and QuPath-0.6^96^. Brain slices were aligned to the Allen Brain atlas using the ABBA plugin for Fiji^97^. Co-localization was quantified using the Cell Detection function in QuPath-0.6.

### Fluorescent *in situ* hybridization (FISH)

Mice were sacrificed with an intraperitoneal injection of 300 mg/kg ketamine and 20 mg/kg xylazine, perfused transcardially with PBS and fixed *in-situ* with 4 % PFA diluted in PBS. Brains were carefully extracted, post-fixed in 4 % PFA for 24 h, followed by 48 h in 30 % sucrose at 4 °C, then embedded in OCT, frozen in dry ice and sliced at 20 µm with a cryostat (Leica catalog # CM1950).

Slices were fixed for 15 min with 4 % PFA, washed with PBS and dehydrated using an ethanol gradient ranging from 25, 50, 75 to 100 % for 5 min each. Fixed slices were then stained with ISH probes for detection of *Fos (*#NM_010234.3) and *Crh* (#NM_205769), which were designed by the manufacturer (Molecular Instruments). Images were obtained using an inverted point laser scanning confocal microscope (Axio Observer, Carl Zeiss AG catalog # LSM980) with a 20x and 40x objective.

## Statistical analysis

Data is depicted as mean ± SD. Statistical significance was calculated as indicated in the respective figure legends using GraphPad Prism 11.

## Data availability

The RNA-seq dataset is available at the European Nucleotide Archive (ENA) at EMBL-EBI under accession number PRJEB113181. The 16S rRNA gene amplicon sequencing data is available at NCBI (https://www.ncbi.nlm.nih.gov/) under the BioProject accession number PRJNA1454696.

## Supplementary figure legends

**Supplementary figure 1:**
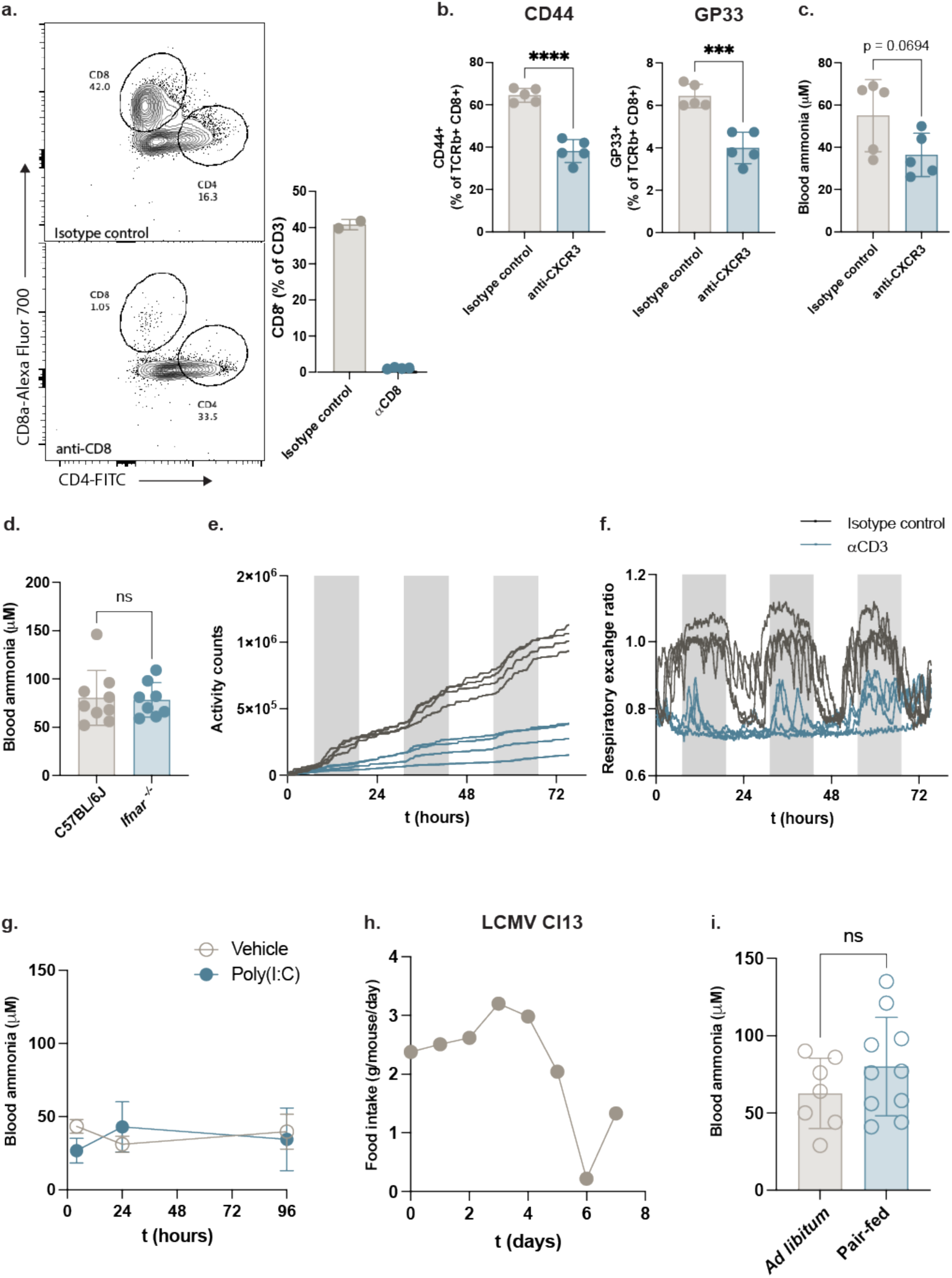
(a) Representative plots of flow cytometric analysis of splenic CD8^+^ T cells 8 days after infection with LCMV Cl13 (left) and respective quantification (right). Mice had been pre-treated with an anti-CD8a depleting antibody or an isotype control. Data shows mean±SD, n=2-4 mice per group representative of two independent experiments. (b) Percentage of CD44^+^ (left) and virus-specific GP33^+^ (right) splenic CD8^+^ T cells 8 days after infection with LCMV Cl13. Mice had been pre-treated with an anti-CXCR3 blocking antibody or an isotype control. Data shows mean±SD, n=5 mice per group.-Statistical analysis: unpaired t-test. (c) Blood ammonia measurements at 8 days post LCMV Cl13 infection in mice treated with an anti-CXCR3 blocking antibody or an isotype control. Data shows mean±SD, n=5 mice per group. Statistical analysis: unpaired t-test. (d) Blood ammonia measurements at 8 days post LCMV Cl13 infection in *Ifnar1^−/−^* mice. Data shows mean±SD, n=8-9 mice per group pooled from two independent experiments. Statistical analysis: unpaired t-test. (e, f) Cumulative activity counts (e) and respiratory exchange ratio (RER) (f) recorded in metabolic cages in C57BL/6J mice treated with 50 μ g anti-CD3 (to induce sterile inflammation) or isotype control. Each dot represents one mouse and shaded areas indicate the dark cycle; n=4 mice per group. (g) Kinetics of blood ammonia in C57BL/6J mice treated with 10 mg/kg poly(I:C) or vehicle. Data represents mean±SD, n=5 mice per group. (h) Average daily food intake of mice infected with LCMV Cl13. (i) Blood ammonia measurements after 8 days of *ad libitum* feeding or pair feeding. Pair-fed mice were given the same amount of food as consumed by LCMV Cl13-infected mice, as shown in (h). Data shows mean±SD, n=7-10 mice per group, pooled from two independent experiments. Statistical analysis: unpaired t-test.

**Supplementary figure 2:**
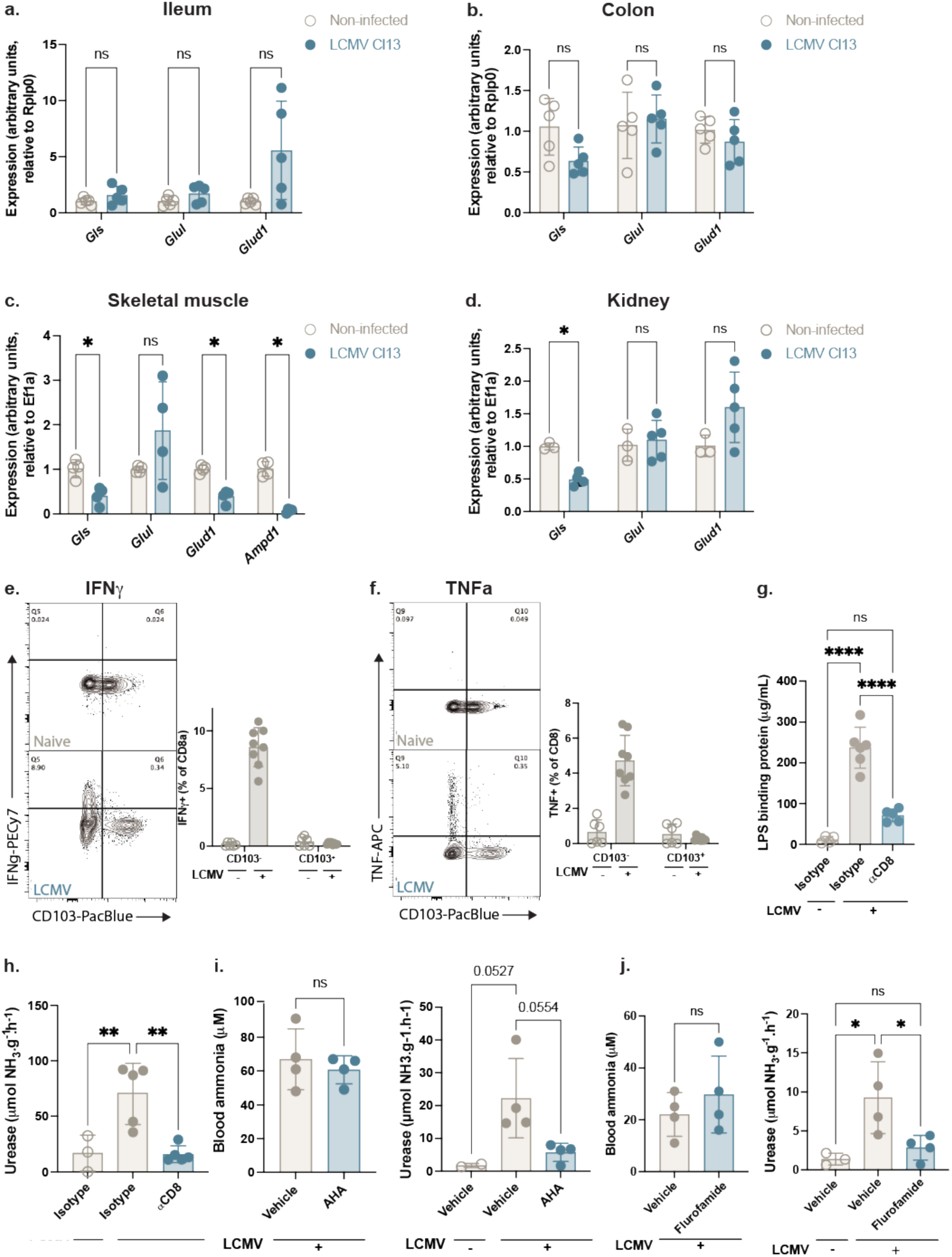
(a-d) Expression analysis of ammonia metabolism genes obtained by quantitative PCR in the ileum (a), colon (b), gastrocnemius muscle (c), and kidney (d) of LCMV-infected mice at 8 dpi. Data shows mean±SD, n=3-5 mice per group. Statistical analysis: multiple unpaired t-tests with Welch correction. (e-f) Flow cytometric analysis of lamina propria lymphocytes (LPL) in the colon of LCMV clone 13 infected mice at 8 dpi and non-infected controls. (e) Percentage of IFN γ producing T cells after re-stimulation with the GP33 viral peptide. (f) Percentage of TNFα producing T cells after re-stimulation with the GP33 viral peptide. Data shows mean±SD, n=6-8 mice per group pooled from two independent experiments. (g) Serum levels of liposaccharide binding protein (LBP) at 8 days post-LCMV Cl13 infection in C57BL/6J pre-treated with an anti-CD8a antibody or an isotype control. Data shows mean±SD, n=4-6 mice per group. Statistical analysis: one-way ANOVA with Tukey multiple comparisons test. (h) Fecal urease activity at 8 days post LCMV Cl13 infection in C57BL/6J pre-treated with an anti-CD8a antibody or an isotype control. Data shows mean±SD, n=3-5 mice per group. Statistical analysis: one-way ANOVA with Tukey multiple comparisons test. (i) Blood ammonia (left) and fecal urease activity (right) at 8 days post LCMV Cl13 infection in mice treated with the urease inhibitor acetohydroxamic acid (AHA) or vehicle control. AHA was administered by oral gavage daily between 5 and 7 dpi. Data shows mean±SD, n=2-4 mice per group. Statistical analysis: one-way ANOVA with Tukey multiple comparisons test. (j) Blood ammonia (left) and fecal urease activity (right) at 8 days post LCMV Cl13 infection in mice treated with the urease inhibitor flurofamide or vehicle control. Flurofamide was administered by intraperitoneal injection at 7 dpi. Data shows mean±SD, n=3-4 mice per group. Statistical analysis: one-way ANOVA with Tukey multiple comparisons test.

**Supplementary figure 3:**
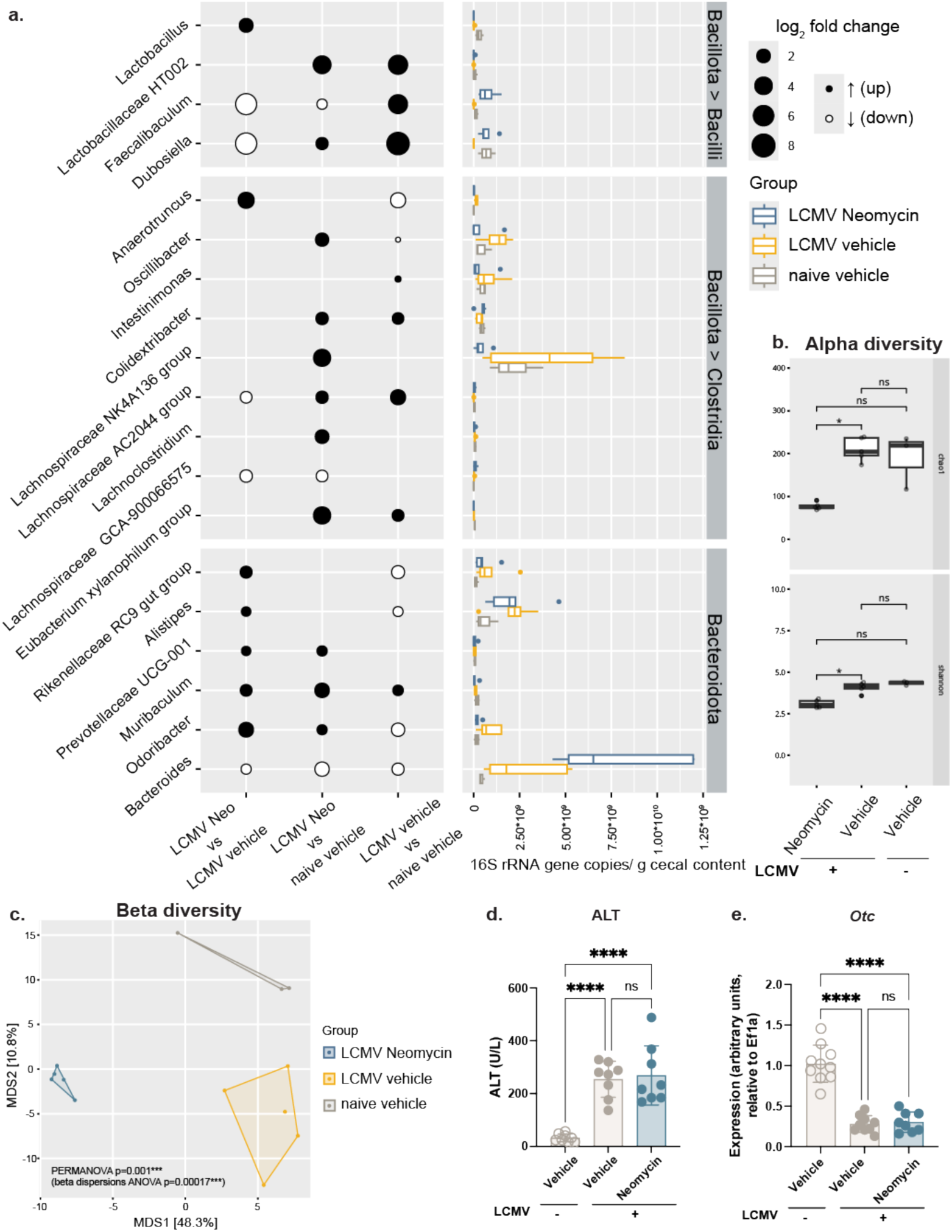
(a) Statistically significant differences between absolute abundances of bacterial genera in the cecum of LCMV Cl13-infected and neomycin-treated mice obtained from 16S rRNA gene amplicon sequencing. n=3-5 mice per group. (b) Alpha diversity index (Chao1 and Shannon) from 16S rRNA gene amplicon sequencing data of cecal samples from LCMV Cl13-infected and neomycin-treated mice. Statistical differences between groups were assessed using pairwise Wilcoxon tests followed by false discovery rate (FDR) correction, n=3-5 mice per group. (c) Beta diversity index (Euclidian distance) from 16S rRNA gene amplicon sequencing data of cecal samples from LCMV Cl13-infected and neomycin-treated mice. Statistical differences between groups were assessed using PERMANOVA, n=3-5 mice per group. (d) Serum alanine transferase (ALT) in mice treated with neomycin as in Figure 2d. Data shows mean±SD, n=8 mice per group pooled from two independent experiments. Statistical analysis: one-way ANOVA with Tukey multiple comparisons test. (e) Expression analysis of *Otc* obtained by quantitative PCR in the liver of mice treated with neomycin as in Figure 2d. Data shows mean±SD, n=8-10 mice per group, pooled from two independent experiments. Statistical analysis: one-way ANOVA with Tukey multiple comparisons test.

**Supplementary figure 4:**
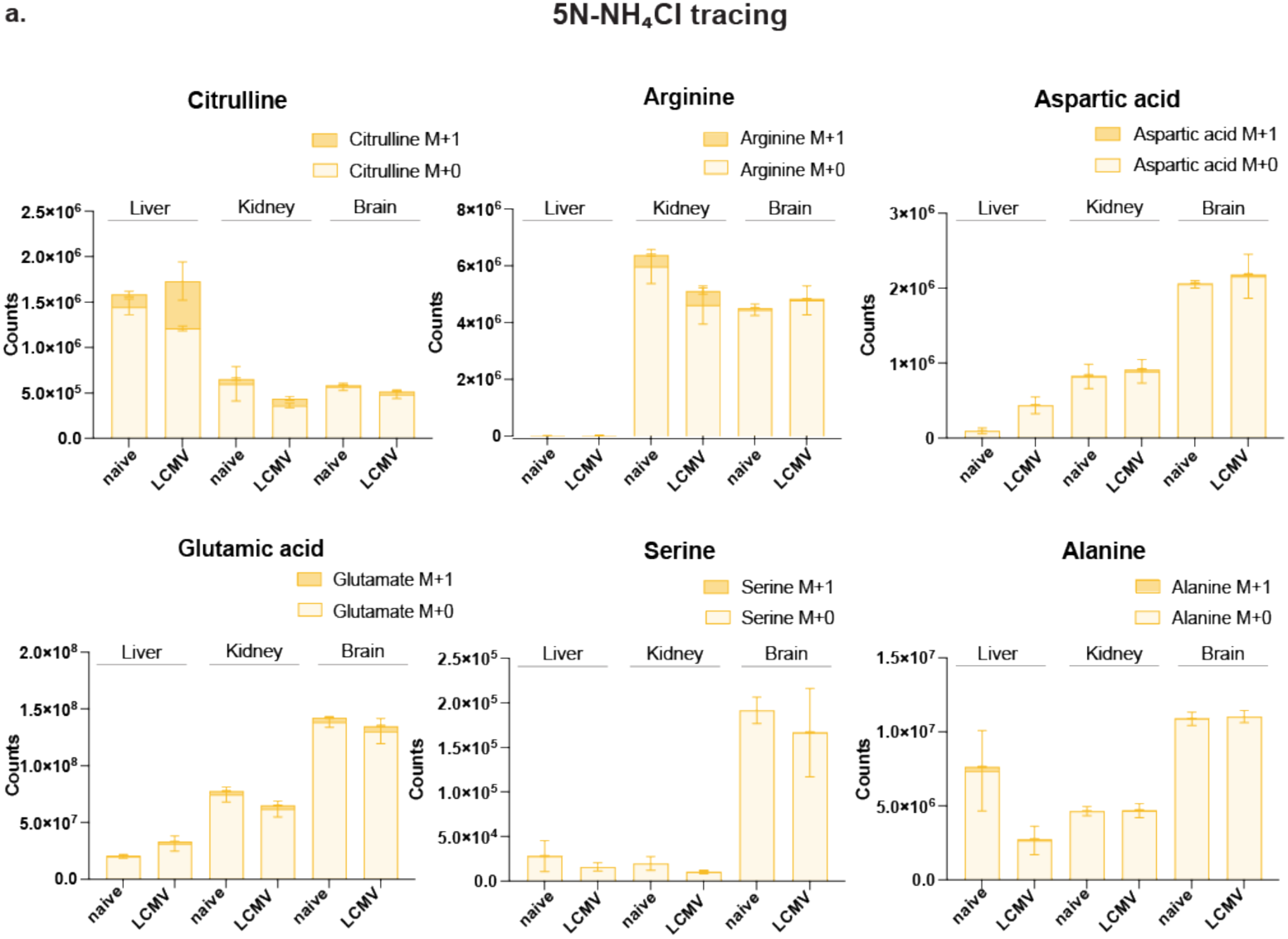
(a) LC-MS quantification of heavy isotope labelled amino acids in different tissues 20 min after intraperitoneal injection of ^15^NH_4_Cl. Data represent mean±SD, n=3 mice per group.

**Supplementary figure 5:**
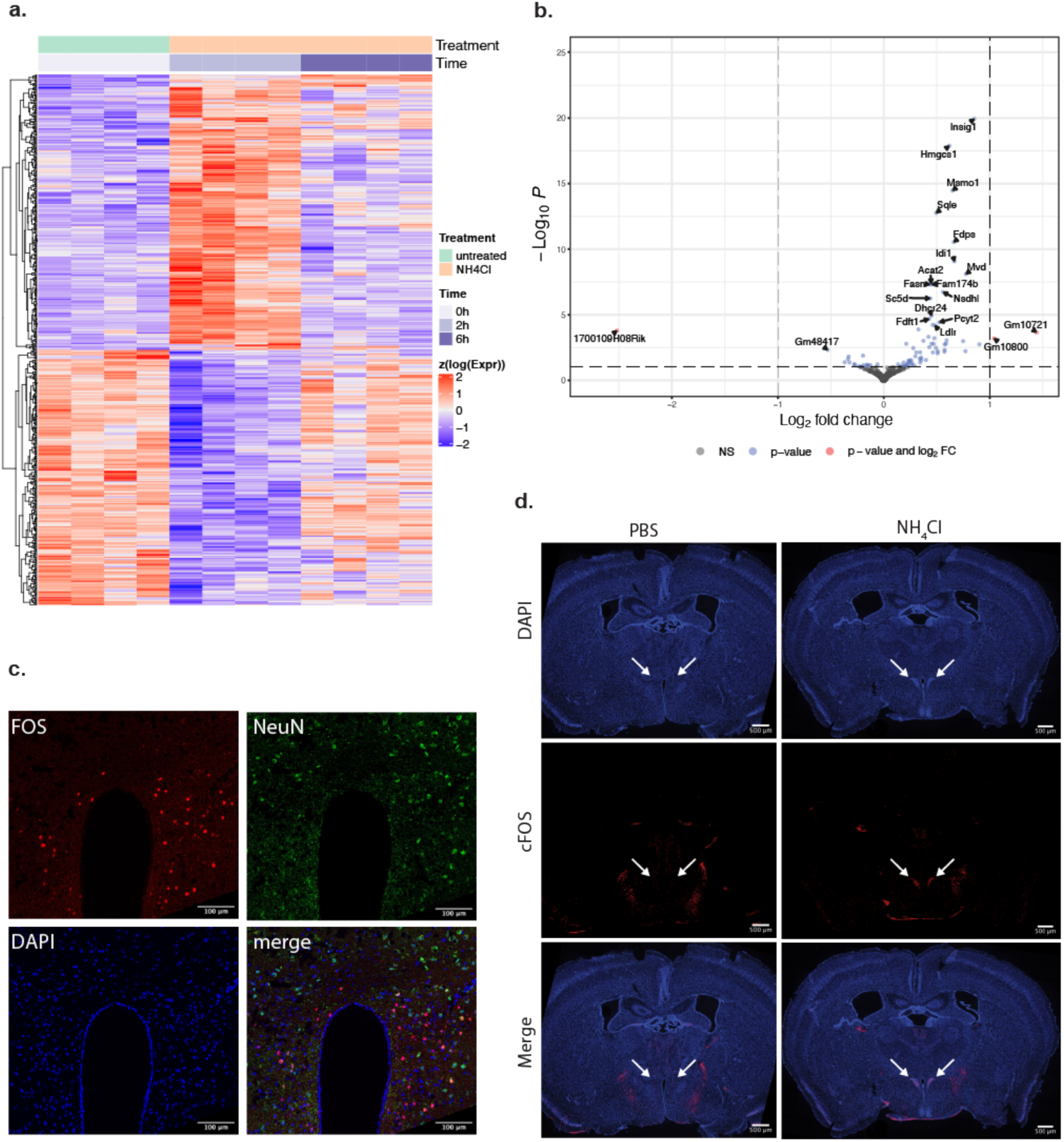
(a) Heatmap of the top 50 most significant genes of mouse embryo cortical neurons incubated with 1 mM NH_4_Cl or PBS for 2 h or 6 h, n=4 mice per group. (b) Volcano plot showing the transcriptional response of mouse embryo cortical neurons incubated with NH_4_Cl or PBS for 6 h. n=4 mice per group. (c) Representative immunofluorescence images showing neuronal (NeuN) and cFOS staining in the PVH of mice 1.5 h after intraperitoneal injection of 200 mg/kg NH_4_Cl or PBS. (d) Representative cFOS immunofluorescence images of a whole brain coronal section 1.5 h after intraperitoneal injection of 200 mg/kg NH_4_Cl or PBS. White arrows indicate the PVH.

**Supplementary figure 6:**
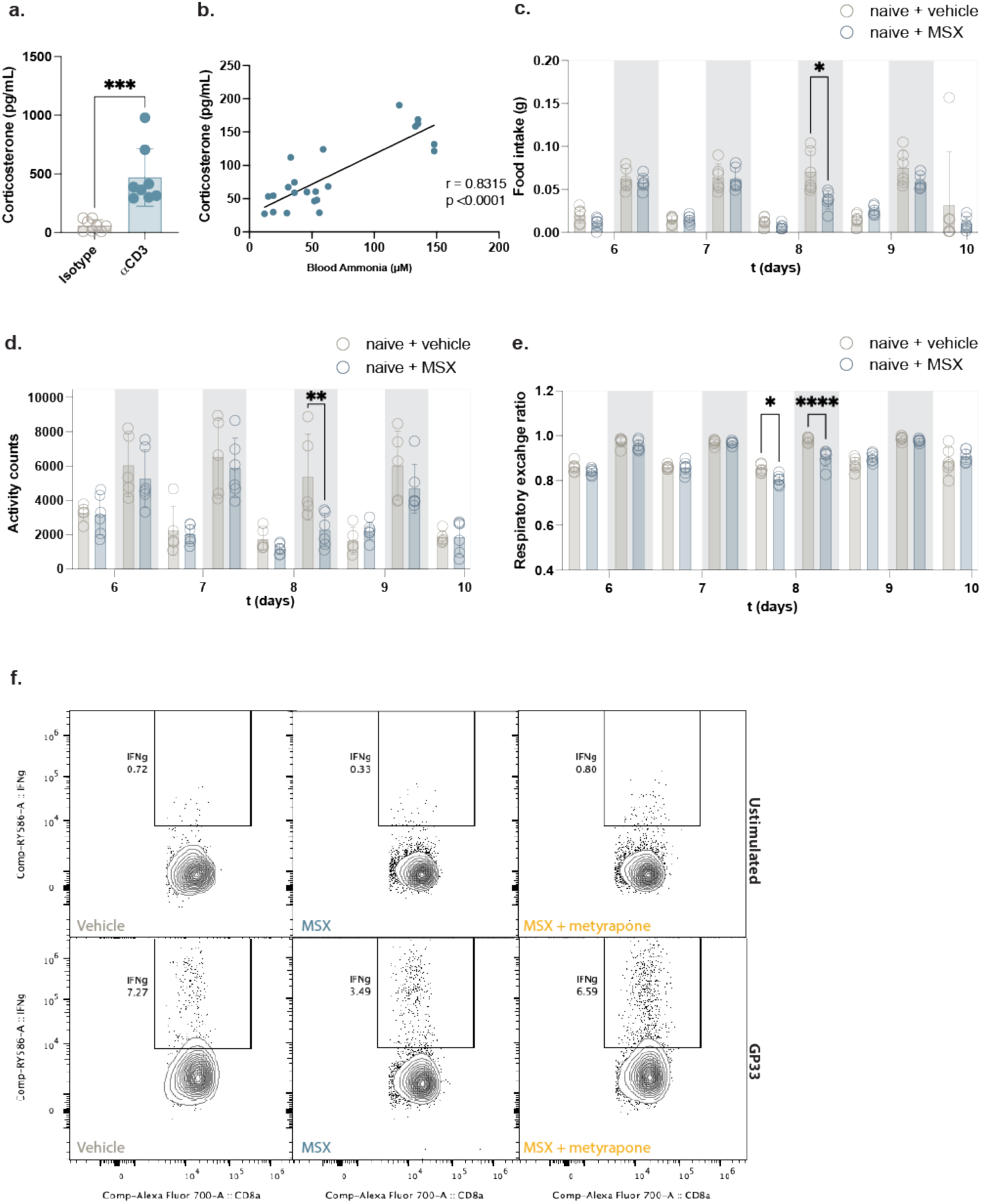
(a) Serum corticosterone measurements 72 h after anti-CD3 injection (to induce sterile inflammation) or isotype control. Data shows mean±SD, n=8 mice per group, pooled from two independent experiments. Statistical analysis: unpaired t-test. (b) Correlation between blood ammonia and serum corticosterone in the same conditions as Figure 5b. Statistical analysis: Pearson correlation test. (c-e) Metabolic cage recordings in non-infected C57BL/6J mice treated with 25 mg/kg MSX: (c) food intake, (d) activity counts, and (e) respiratory exchange ratio (RER). Data shows the average recordings over 15 minute-periods during the active (dark) and rest (light) phases. Shaded areas indicate the dark cycle. n=5-6 mice per group, pooled from two independent experiments. (f) Representative plots of flow cytometric analysis of splenic CD8^+^ T cells 8 days after infection with LCMV Cl13 with MSX and/or metyrapone treatment, as quantified in figure 5i.

## References

1. Wang, A., Luan, H. H. & Medzhitov, R. An evolutionary perspective on immunometabolism. Science (1979). 363, eaar3932 (2019).

2. Troha, K. & Ayres, J. S. Metabolic Adaptations to Infections at the Organismal Level. Trends in Immunology vol. 41 113–125 Preprint at 10.1016/j.it.2019.12.001 (2020).

3. Sammons, M. et al. Brain-body physiology: Local, reflex, and central communication. Cell 187, 5877–5890 (2024).

4. Kraus, A., Buckley, K. M. & Salinas, I. Sensing the world and its dangers: An evolutionary perspective in neuroimmunology. Elife 10, (2021).

5. Eisenberger, N. I., Moieni, M., Inagaki, T. K., Muscatell, K. A. & Irwin, M. R. In Sickness and in Health: The Co-Regulation of Inflammation and Social Behavior. Neuropsychopharmacology 42, 242–253 (2017).

6. Silverman, M. N., Pearce, B. D., Biron, C. A. & Miller, A. H. Immune Modulation of the Hypothalamic-Pituitary-Adrenal (HPA) Axis during Viral Infection. Viral Immunol. 18, 41–78 (2005).

7. Salvador, A. F., de Lima, K. A. & Kipnis, J. Neuromodulation by the immune system: a focus on cytokines. Nat. Rev. Immunol. 1–16 (2021) doi:10.1038/s41577-021-00508-z.

8. Osterhout, J. A. et al. A preoptic neuronal population controls fever and appetite during sickness. Nature 606, 937–944 (2022).

9. Ilanges, A. et al. Brainstem ADCYAP1+ neurons control multiple aspects of sickness behaviour. Nature 609, 761–771 (2022).

10. McCarville, J. L., Chen, G. Y., Cuevas, V. D., Troha, K. & Ayres, J. S. Microbiota Metabolites in Health and Disease. Annu. Rev. Immunol. 38, 147–170 (2020).

11. Labarta-Bajo, L. et al. CD8 T cells drive anorexia, dysbiosis, and blooms of a commensal with immunosuppressive potential after viral infection. Proceedings of the National Academy of Sciences 117, 24998–25007 (2020).

12. Sencio, V. et al. Gut Dysbiosis during Influenza Contributes to Pulmonary Pneumococcal Superinfection through Altered Short-Chain Fatty Acid Production. Cell Rep. 30, 2934–2947.e6 (2020).

13. Uchimura, Y. et al. Antibodies Set Boundaries Limiting Microbial Metabolite Penetration and the Resultant Mammalian Host Response. Immunity 49, 545–559.e5 (2018).

14. Gabanyi, I. et al. Bacterial sensing via neuronal Nod2 regulates appetite and body temperature. Science (1S7S). 376, (2022).

15. Morais, L. H., Schreiber, H. L. & Mazmanian, S. K. The gut microbiota–brain axis in behaviour and brain disorders. Nat. Rev. Microbiol. 19, 241–255 (2021).

16. Wang, R. Gasotransmitters: Growing pains and joys. Trends in Biochemical Sciences vol. 39 227–232 Preprint at 10.1016/j.tibs.2014.03.003 (2014).

17. McCarville, J. L., Chen, G. Y., Cuevas, V. D., Troha, K. & Ayres, J. S. Microbiota Metabolites in Health and Disease. Annual Review of Immunology vol. 38 147–170 Preprint at 10.1146/annurev-immunol-071219-125715 (2020).

18. Wallace, J. L. & Wang, R. Hydrogen sulfide-based therapeutics: exploiting a unique but ubiquitous gasotransmitter. Nat. Rev. Drug Discov. 14, 329–345 (2015).

19. Lundberg, J. O. & Weitzberg, E. Nitric oxide signaling in health and disease. Cell 185, 2853–2878 (2022).

20. Levitt, D. & Levitt, M. A model of blood-ammonia homeostasis based on a quantitative analysis of nitrogen metabolism in the multiple organs involved in the production, catabolism, and excretion of ammonia in humans. Clin. Exp. Gastroenterol. Volume 11, 193–215 (2018).

21. Aldridge, D. R., Tranah, E. J. & Shawcross, D. L. Pathogenesis of hepatic encephalopathy: Role of ammonia and systemic inflammation. Journal of Clinical and Experimental Hepatology vol. 5 S7–S20 Preprint at 10.1016/j.jceh.2014.06.004 (2015).

22. Wang, P. et al. Gut microbiome-derived ammonia modulates stress vulnerability in the host. Nat. Metab. 5, 1986–2001 (2023).

23. Lercher, A. et al. Type I Interferon Signaling Disrupts the Hepatic Urea Cycle and Alters Systemic Metabolism to Suppress T Cell Function. Immunity 51, 1074–1087.e9 (2019).

24. Baazim, H. et al. CD8+ T cells induce cachexia during chronic viral infection. Nat. Immunol. 20, 701–710 (2019).

25. Bangs, D. J. et al. CXCR3 regulates stem and proliferative CD8+ T cells during chronic infection by promoting interactions with DCs in splenic bridging channels. Cell Rep. 38, 110266 (2022).

26. Pontes Ferreira, C., et al. CXCR3 chemokine receptor contributes to specific CD8+ T cell activation by pDC during infection with intracellular pathogens. PLoS Negl. Trop. Dis. 14, e0008414 (2020).

27. Esplugues, E. et al. Control of TH17 cells occurs in the small intestine. Nature 475, 514–518 (2011).

28. Merger, M. et al. Defining the roles of perforin, Fas/FasL, and tumour necrosis factor α in T cell induced mucosal damage in the mouse intestine. Gut 51, 155 (2002).

29. Antonio-Herrera, L. et al. CD8+ T cells regulate the bioenergetic reprogramming of lymphoid organs and the heart during viral infection. bioRxiv 2025.08.20.670307 (2025) doi:10.1101/2025.08.20.670307.

30. Graham, T. E., Turcotte, L. P., Kiens, B. & Richter, E. A. Training and muscle ammonia and amino acid metabolism in humans during prolonged exercise. J. Appl. Physiol. 78, 725–735 (1995).

31. Beura, L. K. et al. Lymphocytic choriomeningitis virus persistence promotes effector-like memory differentiation and enhances mucosal T cell distribution. J. Leukoc. Biol. 97, 217–225 (2015).

32. Labarta-Bajo, L. et al. Type I IFNs and CD8 T cells increase intestinal barrier permeability after chronic viral infection. Journal of Experimental Medicine 217, (2020).

33. Becattini, S. et al. Rapid transcriptional and metabolic adaptation of intestinal microbes to host immune activation. Cell Host Microbe 29, 378–393.e5 (2021).

34. Scott, M. C. et al. Deep profiling deconstructs features associated with memory CD8+ T cell tissue residence. Immunity 58, 162–181.e10 (2025).

35. Wolpert, E., Phillips, Sidney F. & Summerskill, W. H. J. TRANSPORT OF UREA AND AMMONIA PRODUCTION IN THE HUMAN COLON. The Lancet 298, 1387–1390 (1971).

36. Romero-Gómez, M. et al. Intestinal glutaminase activity is increased in liver cirrhosis and correlates with minimal hepatic encephalopathy. J. Hepatol. 41, 49–54 (2004).

37. Jakhar, D., Sarin, S. K. & Kaur, S. Gut microbiota and dynamics of ammonia metabolism in liver disease. npj Gut and Liver 1, 11 (2024).

38. Davila, A.-M. et al. Intestinal luminal nitrogen metabolism: Role of the gut microbiota and consequences for the host. Pharmacol. Res. 68, 95–107 (2013).

39. Li, T.-T. et al. Microbiota metabolism of intestinal amino acids impacts host nutrient homeostasis and physiology. Cell Host Microbe 32, 661–675.e10 (2024).

40. Conrado, L. et al. Anaerobic Conversion of Proteinogenic Amino Acids When Methanogenesis Is Inhibited: Carboxylic Acid Production from Single Amino Acids. Fermentation 10, 237 (2024).

41. Zeng, X. et al. Gut bacterial nutrient preferences quantified in vivo. Cell 185, 3441–3456.e19 (2022).

42. Williamson, G. et al. A two-lane mechanism for selective biological ammonium transport. Elife 9, (2020).

43. Cooper, A. J., McDonald, J. M., Gelbard, A. S., Gledhill, R. F. & Duffy, T. E. The metabolic fate of 13N-labeled ammonia in rat brain. Journal of Biological Chemistry 254, 4982–4992 (1979).

44. Thrane, V. R. et al. Ammonia triggers neuronal disinhibition and seizures by impairing astrocyte potassium buffering. Nat. Med. 19, 1643–1648 (2013).

45. Boutagouga Boudjadja, M., et al. Hypothalamic AgRP neurons exert top-down control on systemic TNF-α release during endotoxemia. Current Biology 32, 4699–4706.e4 (2022).

46. Wu, W.-L. et al. Microbiota regulate social behaviour via stress response neurons in the brain. Nature 595, 409–414 (2021).

47. Tofani, G. S. S. et al. Gut microbiota regulates stress responsivity via the circadian system. Cell Metab. 37, 138–153.e5 (2025).

48. Sommershof, A., Basler, M., Riether, C., Engler, H. & Groettrup, M. Attenuation of the cytotoxic T lymphocyte response to lymphocytic choriomeningitis virus in mice subjected to chronic social stress. Brain Behav. Immun. 25, 340–348 (2011).

49. Acharya, N. et al. Endogenous Glucocorticoid Signaling Regulates CD8+ T Cell Differentiation and Development of Dysfunction in the Tumor Microenvironment. Immunity 53, 658–671.e6 (2020).

50. Souffriau, J. et al. A screening assay for Selective Dimerizing Glucocorticoid Receptor Agonists and Modulators (SEDIGRAM) that are effective against acute inflammation. Sci. Rep. 8, 12894 (2018).

51. Præstholm, S. M., Correia, C. M. & Grøntved, L. Multifaceted Control of GR Signaling and Its Impact on Hepatic Transcriptional Networks and Metabolism. Front. Endocrinol. (Lausanne). 11, (2020).

52. Raabe, W. A. & Onstad, G. R. Ammonia and methionine sulfoximine intoxication. Brain Res. 242, 291–298 (1982).

53. Bernard-Hélary, K., Ardourel, M.-Y., Hévor, T. & Cloix, J.-F. In vivo and in vitro glycogenic effects of methionine sulfoximine are different in two inbred strains of mice. Brain Res. 929, 147–155 (2002).

54. Redford, S. E., Varanasi, S. K., Sanchez, K. K., Thorup, N. R. & Ayres, J. S. CD4+ T cells regulate sickness-induced anorexia and fat wasting during a chronic parasitic infection. Cell Rep. 42, 112814 (2023).

55. Miura, N. et al. Anti-CD3 induces bi-phasic apoptosis in murine intestinal epithelial cells: possible involvement of the Fas/Fas ligand system in different T cell compartments. Int. Immunol. 17, 513–522 (2005).

56. Nouveau, L. et al. Immunological analysis of the murine anti-CD3-induced cytokine release syndrome model and therapeutic efficacy of anti-cytokine antibodies. Eur. J. Immunol. 51, 2074–2085 (2021).

57. Rose, C. F. et al. Hepatic encephalopathy: Novel insights into classification, pathophysiology and therapy. J. Hepatol. 73, 1526–1547 (2020).

58. Zhang, H. et al. Ammonia-induced lysosomal and mitochondrial damage causes cell death of effector CD8+ T cells. Nat. Cell Biol. 26, 1892–1902 (2024).

59. Tang, K. et al. Ammonia detoxification promotes CD8+ T cell memory development by urea and citrulline cycles. Nat. Immunol. 24, 162–173 (2023).

60. Bell, H. N. et al. Microenvironmental ammonia enhances T cell exhaustion in colorectal cancer. Cell Metab. 35, 134–149.e6 (2023).

61. Gu, J. et al. Tumor-produced ammonia is metabolized by regulatory T cells to further impede anti-tumor immunity. Cell 189, 418–434.e24 (2026).

62. Jaschke, N. P. et al. Gut-to-brain signaling restricts dietary protein intake during recovery from catabolic states. Cell 188, 7481–7494.e16 (2025).

63. Li, Y. et al. Ammonia exposure causes the imbalance of the gut-brain axis by altering gene networks associated with oxidative metabolism, inflammation and apoptosis. Ecotoxicol. Environ. Saf. 224, 112668 (2021).

64. Ortega, V. A., Renner, K. J. & Bernier, N. J. Appetite-suppressing effects of ammonia exposure in rainbow trout associated with regional and temporal activation of brain monoaminergic and CRF systems. Journal of Experimental Biology 208, 1855–1866 (2005).

65. Visek, W. J. Ammonia: Its Effects on Biological Systems, Metabolic Hormones, and Reproduction. J. Dairy Sci. 67, 481–498 (1984).

66. Walker, V. Severe hyperammonaemia in adults not explained by liver disease. Annals of Clinical Biochemistry: International Journal of Laboratory Medicine 49, 214–228 (2012).

67. Tetaj, N. et al. Epidemiology, Clinical Presentation and Treatment of Non-Hepatic Hyperammonemia in ICU COVID-19 Patients. J. Clin. Med. 11, 2592 (2022).

68. Bobermin, L. D. & Quincozes-Santos, A. COVID-19 and hyperammonemia: Potential interplay between liver and brain dysfunctions. Brain Behav. Immun. Health 14, 100257 (2021).

69. Honore, P. M. et al. Liver injury without liver failure in COVID-19 patients: how to explain, in some cases, elevated ammonia without hepatic decompensation. Crit. Care 24, 352 (2020).

70. Ruzek, M. C., Pearce, B. D., Miller, A. H. & Biron, C. A. Endogenous Glucocorticoids Protect Against Cytokine-Mediated Lethality During Viral Infection. The Journal of Immunology 162, 3527–3533 (1999).

71. Van den Berghe, G., Téblick, A., Langouche, L. & Gunst, J. The hypothalamus-pituitary-adrenal axis in sepsis- and hyperinflammation-induced critical illness: Gaps in current knowledge and future translational research directions. EBioMedicine 84, 104284 (2022).

72. Su, Y. et al. Multiple early factors anticipate post-acute COVID-19 sequelae. Cell 185, 881–895.e20 (2022).

73. Klein, J. et al. Distinguishing features of long COVID identified through immune profiling. Nature 623, 139–148 (2023).

74. Frank, M. G. et al. SARS-CoV-2 S1 subunit produces a protracted priming of the neuroinflammatory, physiological, and behavioral responses to a remote immune challenge: A role for corticosteroids. Brain Behav. Immun. 121, 87–103 (2024).

75. Turnbull, A. V. & Rivier, C. L. Regulation of the Hypothalamic-Pituitary-Adrenal Axis by Cytokines: Actions and Mechanisms of Action. Physiol. Rev. 79, 1–71 (1999).

76. Foster, J. A., Rinaman, L. & Cryan, J. F. Stress & the gut-brain axis: Regulation by the microbiome. Neurobiol. Stress 7, 124–136 (2017).

77. Müller, U. et al. Functional Role of Type I and Type II Interferons in Antiviral Defense. Science (1979). 264, 1918–1921 (1994).

78. Shinkai, Y. RAG-2-deficient mice lack mature lymphocytes owing to inability to initiate V(D)J rearrangement. Cell 68, 855–867 (1992).

79. Bergthaler, A. et al. Viral replicative capacity is the primary determinant of lymphocytic choriomeningitis virus persistence and immunosuppression. Proceedings of the National Academy of Sciences 107, 21641–21646 (2010).

80. Campbell, C. et al. Bacterial metabolism of bile acids promotes generation of peripheral regulatory T cells. Nature 581, 475–479 (2020).

81. Pjevac, P. et al. An Economical and Flexible Dual Barcoding, Two-Step PCR Approach for Highly Multiplexed Amplicon Sequencing. Front. Microbiol. 12, (2021).

82. Parada, A. E., Needham, D. M. & Fuhrman, J. A. Every base matters: assessing small subunit rRNA primers for marine microbiomes with mock communities, time series and global field samples. Environ. Microbiol. 18, 1403–1414 (2016).

83. Apprill, A., McNally, S., Parsons, R. & Weber, L. Minor revision to V4 region SSU rRNA 806R gene primer greatly increases detection of SAR11 bacterioplankton. Aquatic Microbial Ecology 75, 129–137 (2015).

84. Callahan, B. J. et al. DADA2: High-resolution sample inference from Illumina amplicon data. Nat. Methods 13, 581–583 (2016).

85. Callahan, B. J., Sankaran, K., Fukuyama, J. A., McMurdie, P. J. & Holmes, S. P. Bioconductor Workflow for Microbiome Data Analysis: from raw reads to community analyses. F1000Res. 5, 1492 (2016).

86. Quast, C., et al. The SILVA ribosomal RNA gene database project: improved data processing and web-based tools. Nucleic Acids Res. 41, D590–D596 (2012).

87. Huang, R. et al. TreeSummarizedExperiment: a S4 class for data with hierarchical structure. F1000Res. 9, 1246 (2021).

88. McMurdie, P. J. & Holmes, S. phyloseq: An R Package for Reproducible Interactive Analysis and Graphics of Microbiome Census Data. PLoS One 8, e61217 (2013).

89. Barnett, D., Arts, I. & Penders, J. microViz: an R package for microbiome data visualization and statistics. J. Open Source Softw. 6, 3201 (2021).

90. Fell, C. W. et al. FIBCD is an endocytic GAG receptor associated with a novel neurodevelopmental disorder. EMBO Mol. Med. 14, (2022).

91. Patro, R., Duggal, G., Love, M. I., Irizarry, R. A. & Kingsford, C. Salmon provides fast and bias-aware quantification of transcript expression. Nat. Methods 14, 417–419 (2017).

92. Stolarczyk, M., Reuter, V. P., Smith, J. P., Magee, N. E. & Sheffield, N. C. Refgenie: a reference genome resource manager. Gigascience 9, (2020).

93. Soneson, C., Love, M. I. & Robinson, M. D. Differential analyses for RNA-seq: transcript-level estimates improve gene-level inferences. F1000Res. 4, 1521 (2016).

94. Durinck, S., Spellman, P. T., Birney, E. & Huber, W. Mapping identifiers for the integration of genomic datasets with the R/Bioconductor package biomaRt. Nat. Protoc. 4, 1184–1191 (2009).

95. Love, M. I., Huber, W. & Anders, S. Moderated estimation of fold change and dispersion for RNA-seq data with DESeq2. Genome Biol. 15, 550 (2014).

96. Bankhead, P. et al. QuPath: Open source software for digital pathology image analysis. Sci. Rep. 7, 16878 (2017).

97. Chiaruttini, N. et al. ABBA+BraiAn, an integrated suite for whole-brain mapping, reveals brain-wide differences in immediate-early genes induction upon learning. Cell Rep. 44, 115876 (2025).

